# A heterogeneous population code at the first synapse of vision

**DOI:** 10.1101/2024.05.03.592379

**Authors:** Tessa Herzog, Takeshi Yoshimatsu, Jose Moya-Diaz, Ben James, Leon Lagnado, Tom Baden

**Affiliations:** School of Life Sciences, University of Sussex, Brighton BN19QG, UK; Washington University in St Louis, USA; Northwestern University, Chicago, USA

## Abstract

Vision begins when photoreceptors convert light fluctuations into temporal patterns of glutamate release that drive the retinal network. The input-output relation at this first stage has not been systematically measured *in vivo* so it is not known how it operates across a photoreceptor population. Using kHz-rate glutamate imaging in zebrafish, we find that individual red cones encode visual stimuli with high reliability and time-precision, but routinely vary in sensitivity to luminance, contrast, and frequency across the population. Variations in input-output relations are generated by feedback from the horizontal cell network that effectively decorrelate feature representation. A model capturing how zebrafish sample their visual environment indicates that this heterogeneity expands the dynamic range of the retina to improve the coding of natural scenes. Moreover, we find that different kinetic release components are used to encode distinct stimulus features in parallel: sustained release linearly encodes low amplitude light and dark contrasts, but transient release encodes large amplitude dark contrasts. Together, this study reveals an unprecedented degree of functional heterogeneity within same-type photoreceptors and illustrates how separation of different visual features begins in the first synapse in vision.

## INTRODUCTION

The light-encoding properties of photoreceptors underpin the performance limits of vision^1,2^. While we have a detailed understanding of the processes that convert light into a photocurrent^3,4^, we have much less understanding of the output signal by which photoreceptors drive the retinal circuit - the synaptic release of glutamate^5–7^. Here we use larval zebrafish to make an *in vivo* investigation of the way in which cone photoreceptors encode visual stimuli.

Spatial correlations in natural visual scenes^8,9^ cause most stimuli to be encoded by the simultaneous activity of several cones. It is therefore important to understand how the input-output relation varies across a population of same-type photoreceptors^10,11^. A similar relation across the population would indicate a high degree of redundancy in the signal transmitted at the first stage of vision but it is unclear if this is the case^7,10,12^ because individual cones can be re-tuned by cone-intrinsic factors^6,7,13^, their surrounding circuitry^6,14,15^ and/or the local neuromodulator environment^16^. We have investigated how a population of cones encode visual stimuli by monitoring glutamate release from synaptic pedicles *in vivo*. We focus on ‘ancestral red cones’ (for definitions see Ref ^17^), which likely represent the ancestral general-purpose greyscale system of the vertebrate eye^17–19^. More than 90% of cones in the human eye, comprising both ‘red/L’ and ‘green/M’ variants^20^, are of this highly conserved photoreceptor type^21–23^.

We find that while individual red cones generate synaptic outputs that are exceptionally reliable and precise in time, the population as a whole is heterogenous in terms of sensitivity to luminance and contrast. For example, while some cones exhibited approximately linear contrast-response functions, others were strongly rectifying, signalling negative contrasts more strongly than positive. Similarly, while glutamate release rates from some cones reliably followed temporal contrast up to 20 Hz, other cones ceased to respond above 8 Hz. Blocking inhibitory feedback from horizontal cells causes all red cones to transmit signals at lower frequency and become strongly biased to negative contrasts, demonstrating that the outer retinal circuitry plays a key role in determining the input-output relation of the first neurons in vision. A model of bipolar cells summing inputs from cones as zebrafish sample their visual environment indicates that the heterogeneity in cone signals expands the dynamic range of the retina to improve the coding of natural scenes.

## RESULTS

### Isolating the synaptic output of individual red cones

Cones drive the retinal circuit through synaptic pedicles that are invaginated by post-synaptic bipolar cells and horizontal cells to create extracellular compartments tightly sealed against the surrounding circuit^5,24^ (Fig. 1A,B). In larval zebrafish, there is only one such invagination per cone^14^, and we exploited this anatomical arrangement to use the dendritic tips of horizontal cells expressing the fluorescent glutamate sensor SFiGluSnFR^25^ as “glutamate antennae”. A spatially isolated SFiGluSnFR “hotspot” therefore provided a glutamate read-out from an individual cone pedicle^6,7^, each typically comprising some 2-3 ribbons with a total surface area of 0.2-0.5 µm^2^ (Supplemental Figure S1). Applying line scans at 1 kHz typically allowed us to sample the outputs of 2-5 cone pedicles simultaneously (Supplemental Video V1).

**Figure 1.**
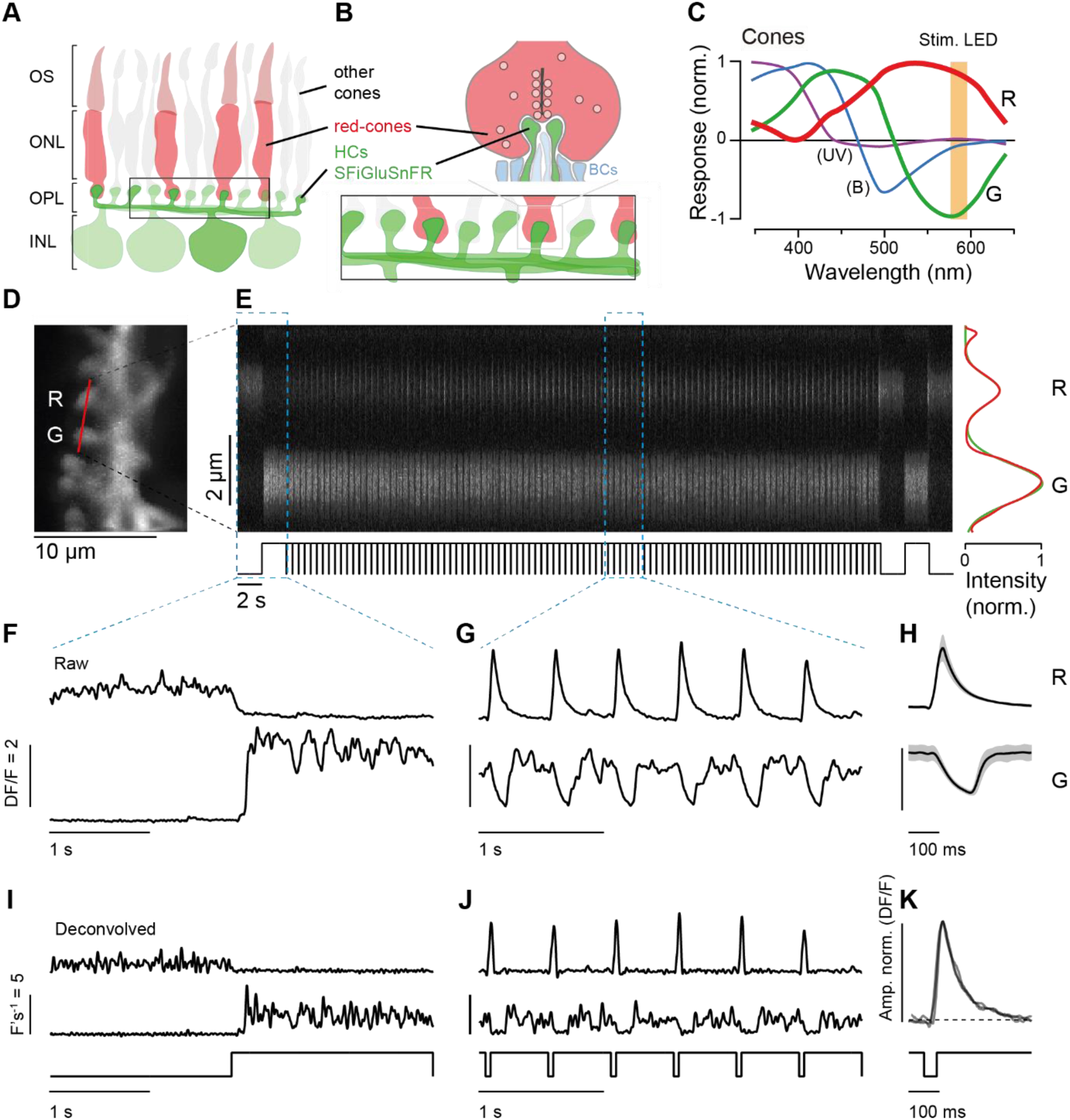
In vivo monitoring of light driven glutamate release from zebrafish ancestral red cones. **A,B**, Dendritic tips of horizontal cells invaginate the synaptic pedicles of cones, and we used SFiGluSnFR-expression in HCs as postsynaptic glutamate antennas of cone release (see also Ref^6^). **C**, In vivo spectral tuning of larval zebrafish cone types (‘red, green, blue, UV’) with 590 nm stimulus wavelength overlaid (modified from Ref ^27^). **D**, Maximum intensity projection of the field of view from a typical two-photon recording, focused on a portion of the nasal outer retina of a 6 dpf larval zebrafish expressing SFiGluSnFR in HCs. Red line indicates approximate positioning of line scan in (E). **E**, Kymograph of the line scan indicated in (D) during widefield visual stimulation. Right: the fluorescence signals over time from two neighbouring cone pedicles (labelled ‘R’ and ‘G’) were extracted based on the two Gaussians that best approximated their spatial profile (based on Ref^26^). **F,G**, Fluorescence traces from (E) for two different parts of the stimulus sequence, as indicated, and **H**, the corresponding mean responses to 98 off-steps of light (± sd). **I,J**, Deconvolved versions of fluorescence traces shown in (F,G, Methods). **K**, Superposition of three amplitude-normalised responses from the red cone in (E) illustrates their highly stereotyped time-courses.

To isolate responses of red cones, we used wide-field stimuli of amber light (∼590 nm) because in larval zebrafish only red cones exhibit excitatory responses to light decrements at this wavelength^14^ (Fig. 1C). Green cones are also sensitive to amber light, but they are red-opponent and therefore exhibit *inhibitory* responses to the same light decrements^14^. The reliability of this strategy for identifying red cones was confirmed by co-labelling red cones with tdTomato (Supplemental Figure S2).

Examples of synaptic responses are shown in Fig. 1D-H. At the start of this experiment, a light step was applied, bright enough to reduce the fluorescence signal in the pedicle labelled R to close to background fluorescence levels while also suppressing the noise caused by vesicle fusion in the dark, indicating complete, or almost complete, suppression of glutamate release (Fig. 1F). From this bright background, off-steps were applied by turning off the light for 40 ms. This pedicle was identified as belonging to an ancestral red cone because it exhibited large amplitude glutamate transients in responses to the off-steps (Figs. 1G-H). In contrast, the neighbouring pedicle labelled “G” exhibited transient *suppressions* of glutamate release, indicating that it belonged to a green cone. Beyond these two basic response types, other pedicles did not generate reliable SFiGluSnFR signals, indicating that they belonged to blue and UV cones (not shown).

A notable feature of the glutamate transients generated by red cone pedicles was that when the amplitude varied, the shape did not (Fig. 1K). This property allowed us to assess relative changes in the rate of glutamate release by deconvolution of the SFiGluSnFR signal with the temporal kernel provided by the average response to a brief off-steps (Figs. 1I,J). A similar approach has been used to detect individual vesicle release events at the synapse of retinal bipolar cells^26^. In cones, however, it was not possible to reliably disambiguate the signals from single vesicles. Instead, we interpret our cone data as reflecting the summed signals from multiple vesicles released from multiple ribbons^7^. Below, we use this technique to investigate how cone synapses encode light intensity and contrast.

### Variable sensitivity to light at the synaptic output of individual cones

The usual approach to investigating the light sensitivity of photoreceptors, for instance when measuring photocurrents, would be to apply brief flashes of fixed duration but different intensities on top of a dark background^28^. We did not use this approach because the dark signal at the cone synapse is noisy (e.g. Fig. 1F,I) and the response to light is a *decrease* in the SFiGluSnFR signal caused by an interruption in the continuous release of vesicles. Measuring a decreasing signal from a high and noisy baseline degrades the signal to noise ratio compared to measuring an increasing signal from a low and stable baseline. We therefore carried out these measurements using off-steps, where the strength of the stimulus was the number of photons “missing” when the light was turned off, in turn directly proportional to the duration. A sequence of off-steps, 5-100 ms in duration were presented in a pseudorandom order. This revealed that different red cones displayed very different sensitivities to light, as illustrated by the two nearby pedicles in Fig. 2A-D (see also Supplemental Video V2). While cone 1 reliably responded to 10 ms flashes and saturated at 20 ms, cone 2 only started to respond to flashes of ∼18 ms and this response did not saturate until the flash duration was 50 ms (Fig. 2C,D).

**Figure 2.**
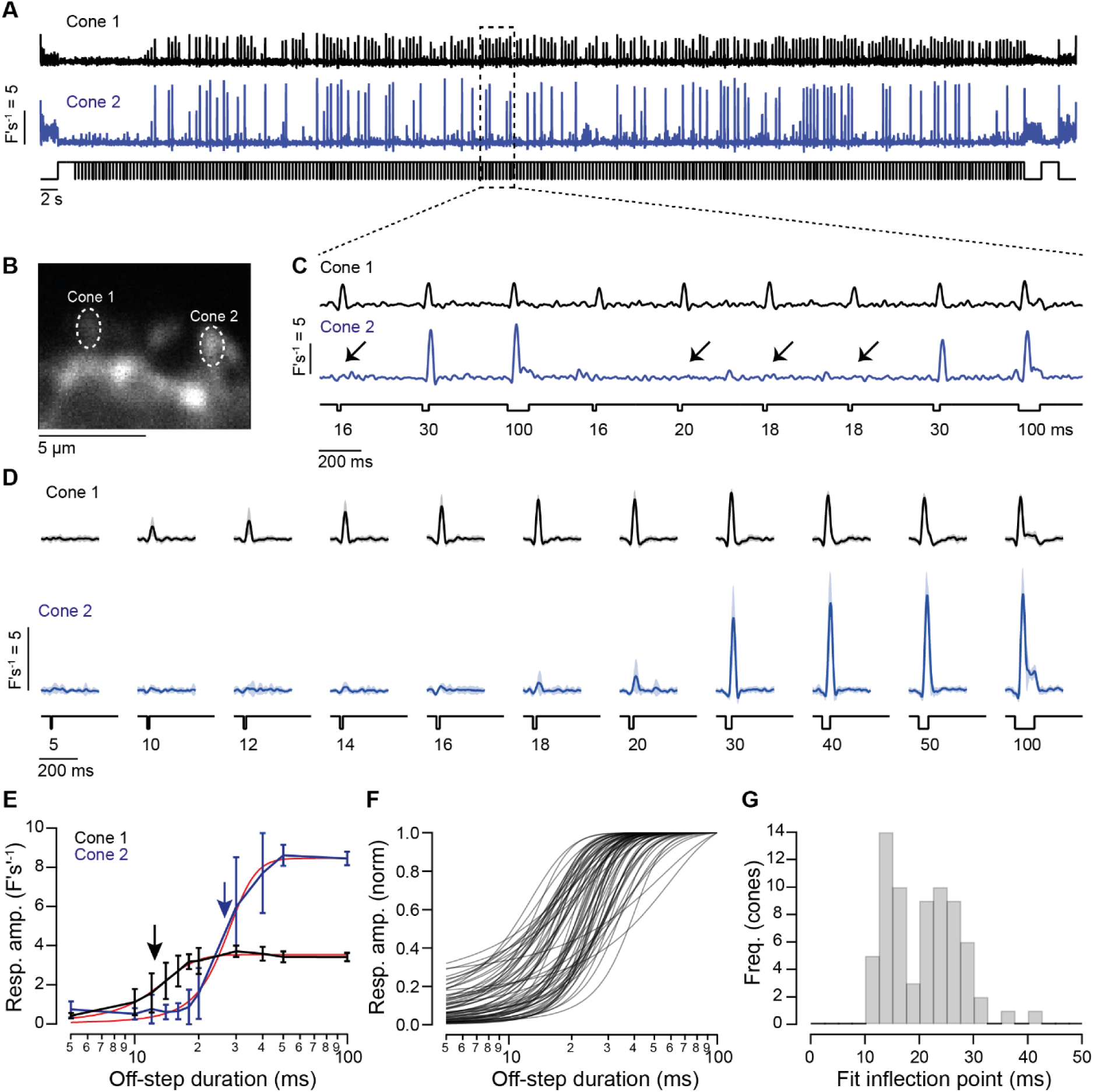
Distinct off-step-sensitivities in neighbouring red cones in vivo. **A-D**, Simultaneously recorded glutamate release (A,C,D) from two neighbouring cones (B) in response to a pseudorandom sequence of 100% contrast off-steps of varying duration (5 to 100 ms). A and C show raw deconvolved signals at two timescales as example cones from A-D, with sigmoidal fits (red). **F**, Amplitude-normalised fits of contrast-response functions of n = 70 cones from 12 fish and **G**, summary of their corresponding inflection points d_1/2_ (see also arrowheads in E, Methods). Hartigan Dip test statistic = 0.046, indicating bimodality. However, fit inflection points strongly correlated with other metrics of heterogeneity (Supplemental Figure S3), and Principal Component Analysis across these metrics did not reveal robust substructure in the overall population of red-cone responses.

The relationship between flash duration (d) and relative rate of glutamate release (R) could be described as a sigmoid function,

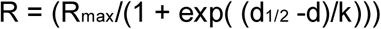

where R_max_ is the saturating response, d_1/2_ is the off-stepduration generating the half-maximal response and k is a constant (Fig. 2E). The stimulus-response functions for cone 1 and 2 differed both in terms of R_max,_ reflecting synaptic gain, and d_1/2_, reflecting light sensitivity. Qualitatively similar differences were routinely observed in simultaneously recorded neighbouring red cones, indicating that response heterogeneity reflected biological rather than experimental variation. A series of fits to the stimulus-response function of 70 cones from 12 fish, normalized to the maximum response, is shown in Fig. 2F and the distribution of d_1/2_ in Fig. 2G. The d_1/2_ value ranged from 10.44 to 40.86 ms, and the mean value of d_1/2_ was 20.83 ± 6.64 ms (± sd). These results demonstrate a substantial degree of response heterogeneity across red cones *in vivo*.

### High reliability and temporal precision of the synaptic signal

A key property of any neural signal is its reliability. Neural information is degraded by noise that causes responses to vary when a stimulus is repeated, and synapses are a major source of such noise because of the stochasticity of the presynaptic processes that control vesicle fusion^2,29,30^. At many central synapses these processes obey Poisson statistics^31^ with a coefficient of variation (CV = sd/mean) of 1. Although it has sometimes been assumed that ribbon synapses of cones also obey Poisson statistics^32^, more recent work indicates that this is not the case in rods^33^.

To explore the reliability of the cone output *in vivo*, we delivered many off-steps (usually 98) of a fixed duration (40 ms) (Fig. 3A). For *individual* red cones, the amplitudes of responses to this repeated stimulus were consistently found to be normally distributed (Fig. 3B). The scatter plot in Fig. 3C shows the relation between the standard deviation and mean of the responses output from 32 cones: in all cases the CV was far below 1 (dashed line) and averaged 0.13 ± 0.05 F’s^−1^ (mean ± sd; Fig. 3D). Re-expressing this metric as the signal-to-noise ratio (SNR = mean^2^/variance) yields an average of 93 for the output of one pedicle. The SNR will depend on the square root of the number of ribbons if these are affected by independent sources of noise. With this assumption and taking an average of two ribbons per pedicle we can place a *lower* limit of SNR = 66 for a single ribbon synapse.

**Figure 3.**
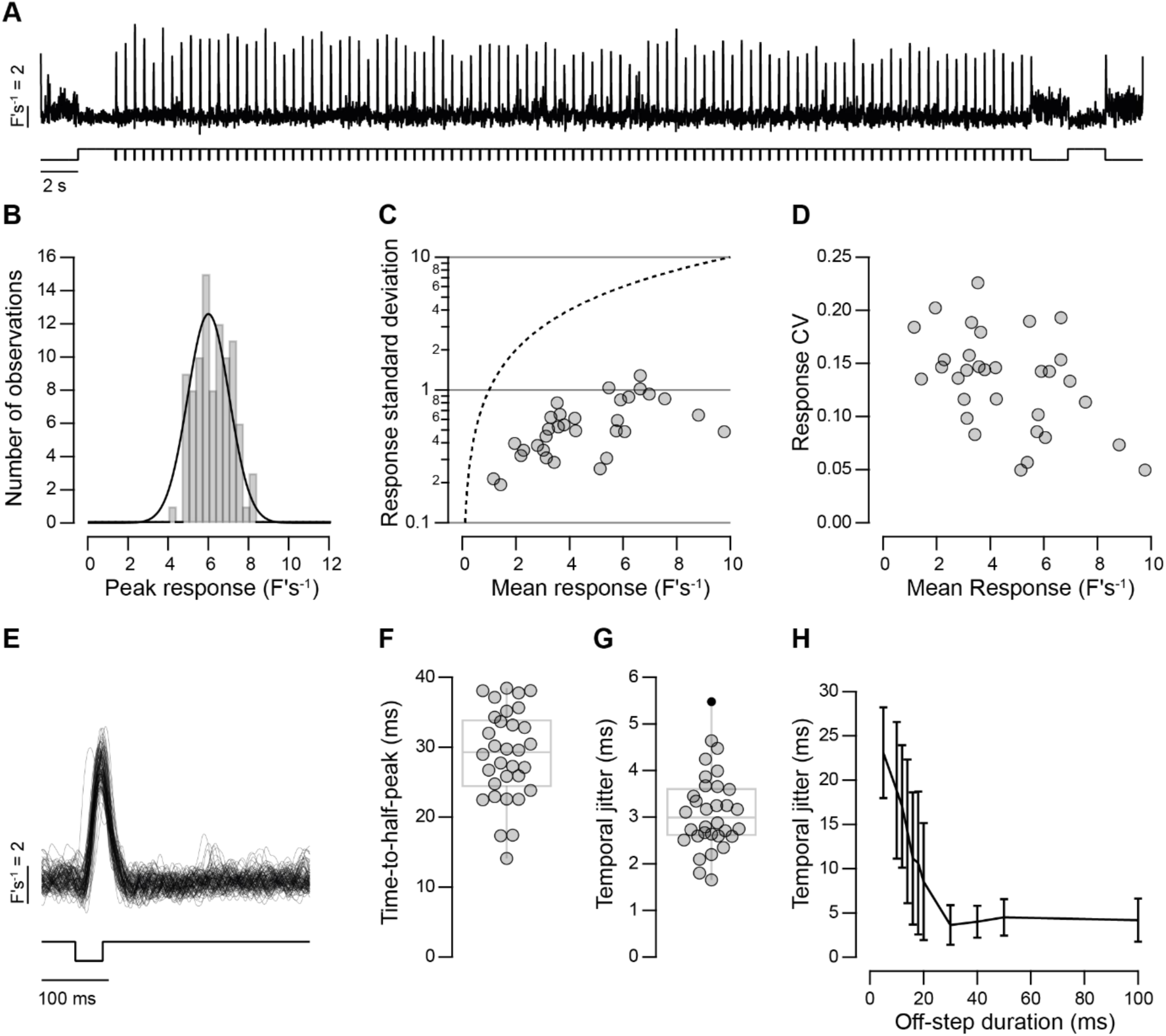
The cone output is reliable and temporally precise. **A,B**, Glutamate responses of one example cone to 98 identical 40 ms duration off-steps and (B) distribution of response amplitudes with Gaussian fit superimposed (mean = 6.2 ± 0.9 (± sd)). **C**, Relationships of mean versus standard deviation of response amplitudes from n = 32 cones in 13 fish systematically fall below the equivalence line where the mean equals the standard deviation, as would be expected from a Poisson release process (dashed). **D**, The coefficient of variation of the data shown in C (Pearson correlation, r = -0.45 p < 0.05). **E-G**, Overlay of all 98 responses from (A) demonstrates a high degree of temporal precision, here quantified as time-to-half-peak (F), and temporal jitter (G, standard deviation of time-to-half-peak). **H**, Temporal jitter was stable for stimulus durations above 30 ms (n = 70 cones, 12 fish).

Responses of sensory neurons vary in their timing as well as their amplitude. To investigate this aspect of the signal transmitted from cones we began by measuring the time-to-half peak (t_1/2_) of responses to off-steps, as shown in Fig. 3E. The mean value of t_1/2_ after a 40 ms off-step was 28.9 ± 6.5 ms across a sample of 32 red cones (Fig. 3F). The temporal precision of these responses was measured as the standard deviation in t_1/2_ at each cone output (Fig. 3G). Some cones displayed temporal jitter in the order of 1-2 ms, and the average was 3.1 ± 0.8 ms. Similar levels of temporal precision were observed for flashes of 30 ms and longer, but jitter increased as off-steps became shorter (Fig. 3H). A temporal precision in the order of a few milliseconds can also be observed in the responses of bipolar cells and ganglion cells responding to stimuli of high contrast^34,35^.

Together, the results in Figs. 1-3 demonstrate that while the ribbon synapses of ancestral red cones *in vivo* transmit the first visual signal with extreme reliability and temporal precision (Fig. 3), the luminance sensitivity varies substantially across the population (Fig. 2).

### Variations in sensitivity to contrast

Natural scenes rarely contain an off-step to darkness. The major task of the cone array is to encode continuous fluctuations in light intensity around an intermediate background (Fig. 4), occurring at different temporal frequencies (Fig. 5). The fact that intensity varies up and down is critical because neurons are rarely linear – changes in opposite directions are often encoded unequally. Synapses are key sites for such rectification. We therefore asked how far the first synapse in vision rectifies and found that the answer depended on timescale (Fig. 4). On short timescales, red cones encode dark-transitions, but not light-transitions, with a transient burst of glutamate release. However, after this transient phase a more sustained component of release allowed some cones to linearly represent both positive and negative contrasts. We reach the above conclusions after presenting 0.5 s steps from a photopic background (± 10-100% contrast, 1 Hz, 50% duty cycle). Fig. 4A-C shows representative responses and their basic quantification from three red cones. Consistent with previous measurements of ribbon-mediated release at photoreceptor synapses^7,36^, responses consistently displayed both transient and sustained components (Fig. 4D-F). Cone 1 responded to both negative and positive contrasts up to 50% in an approximately linear manner, cone 2 only responded to strong negative contrasts, and cone 3 mainly responded to positive contrasts. A range of contrast-response functions were observed in our sample of 97 cones. The most consistent were generated by the transient output and this was always biased to negative contrasts (Off; Fig. 4E). The sustained component was more linear around low contrasts before saturating in some cases (On; Fig. 4F). To quantify the linearity of each response component, we computed a “dark-light index” (DLI) following ref^10^:

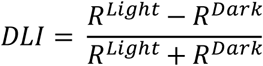

**Figure 4.**
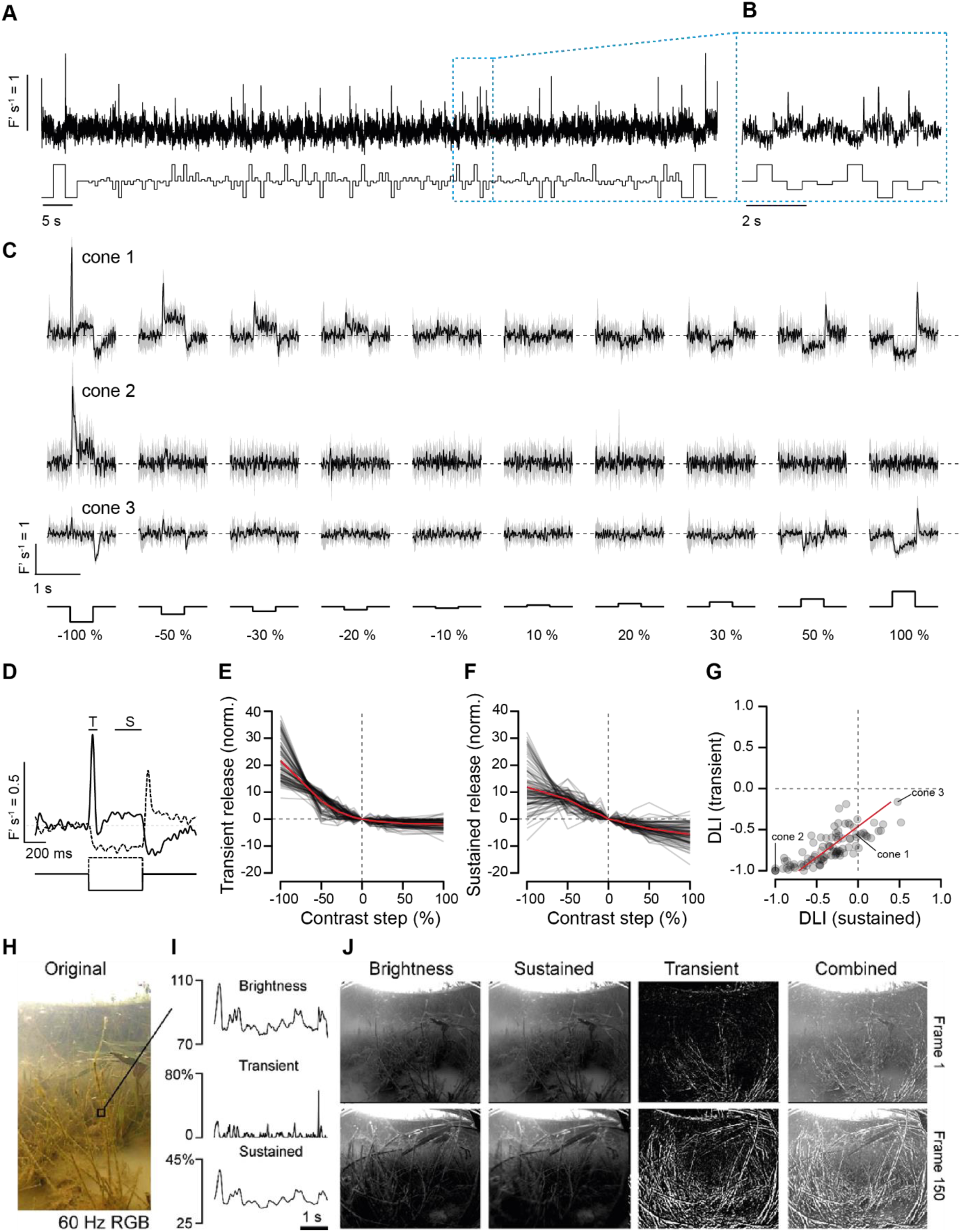
Differential encoding of positive and negative contrasts. **A**,**B**, Glutamate release from one example cone to a pseudorandom sequence of 500 ms light and off-steps from an intermediate baseline shown at two different timescales as indicated. **C**, cone 1, as (A,B), arranged by contrast (mean ± sd (shading)), alongside two further cones processed in the same way (cones 2 and 3). Note that the three example cones exhibit different types of light/dark biases. **D**, Expansion and superposition of cone 1’s mean 100% positive and negative contrast responses as shown in C (dashed and solid lines, respectively) with transient (T) and sustained (S) response components annotated. **E,F,** Contrast response functions of transient (E) and sustained (F) release components of n = 97 red cones from 30 fish with mean superimposed (red). **G**, Distribution of each cone’s dark-light-index (DLI, Methods) of transient and sustained components and linear fit (red). **H-J**, A simple model illustrates how slow linear (here: “sustained”) and fast non-linear (here “transient”) encoding can highlight very different aspects of the same natural scenes (Methods), here showing translational movement through a typical shallow underwater scene from a zebrafish natural habitat in India (from Ref^37^). Each RGB-input-pixel (H) was converted into an 8-bit brightness signal over time (I, top), and then filtered to mimic a strongly dark -biased transient cone response (I, middle), or a linear sustained response (I, bottom). The results are summarised in (J) for two movie frames as indicated. Note that the sustained representation recapitulates most detail in the input, while the transient representation separates out the high-contrast dark edges, which are largely concentrated in the foreground. See also Supplemental Video V2.

**Figure 5.**
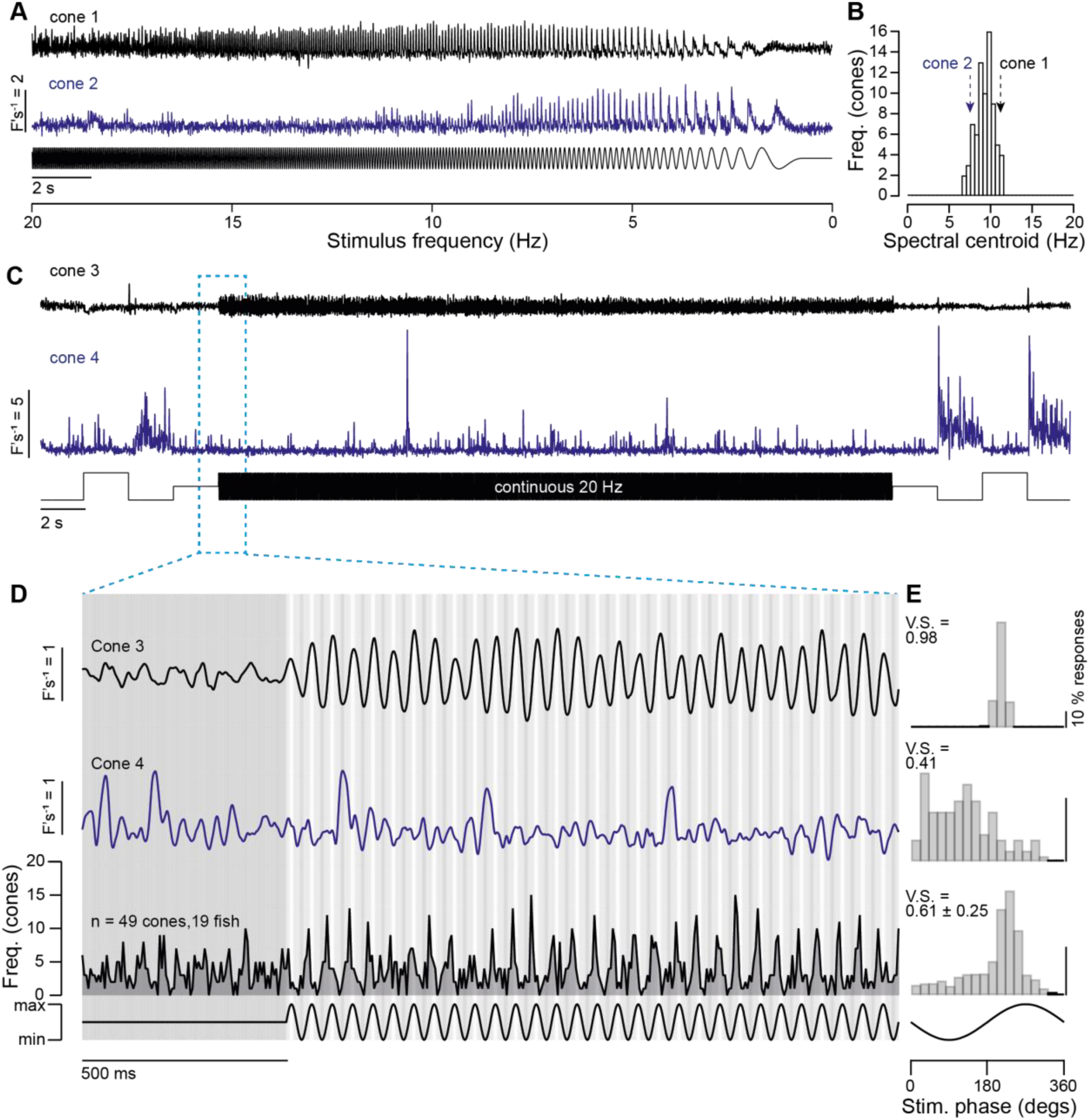
Heterogeneous encoding of temporal contrast. **A**, Release from two example cones driven by a chirp stimulus, decelerating over 20 s from 20 Hz at 100% contrast. **B**, Histogram of spectral centroids of glutamate responses to the stimulus shown in (A) from n = 49 red cones, 18 fish. **C,D**, release from two further examples cones in response to continuous 20 Hz sinusoidal modulation at 100% contrast, with an expansion of the traces shown in (D), and population histogram of detected release events from the population data (bottom, n = 49 cones, 18 fish). **E**, Distributions of detected responses during the stimulus phase for cone 3 (top), cone 4 (middle), and for the population data (bottom) which have a vector strength of 0.98, 0.41, and 0.61, respectively.

where *R^light^* and *R^dark^* are the corresponding areas under the curve for the contrast response functions of transient or sustained responses (Methods). Accordingly, DLI ranged from -1 (responding exclusively to negative contrasts) to 1 (exclusively positive contrasts), while 0 indicated a balanced (linear) contrast-dependence. This revealed that both transient and sustained DLI varied greatly across cones, but while the sustained component spanned almost the entire possible coding range, including strongly positively rectified cones (e.g. cone 3), the transient components were always dark-biased (Fig. 4G). Nevertheless, across cones, the DLI of transient and sustained components covaried strongly (Pearson’s correlation coefficient; *r* = 0.78, *p* < 0.001), suggesting a single mechanism controlling rectification. Further strong positive correlations were found between various additional metrics of heterogeneity (Supplemental Figure S3A-E). Moreover, red-cone heterogeneity was robustly observed in different eye regions, including nasal, dorsal, and the acute zone, with no significant difference in the distribution of DLI values between retinal regions (Supplemental Figure S4A,C).

What might be the consequences of such kinetically unbalanced encoding of positive and negative contrasts? To explore this question, we used stimuli based on natural scenes and a simple model that captured these basic properties of cone release. Based on an underwater video of translatory optic flow taken in the natural habitat of zebrafish (from ref ^37^) (Fig. 4H), we processed each pixel’s brightness sequence over time to mimic representative transient and sustained release components of cones, respectively (Supplemental Video V3, Fig. 4I,J, Methods): Specifically, we differentiated and then negatively rectified each brightness sequence to mimic fast and dark-biased transient release, but amplitude-compressed and time-smoothed each brightness sequence to mimic sustained and linear release. This illustrated how an approximately linear sustained component would yield a time-blurred but otherwise relatively faithful representation of the scene (Fig. 4J, compare first two sets of panels). By contrast, the transient component delivered a very different, parallel representation that was strongly biased to features creating negative contrast in the foreground (Fig. 4J, 3^rd^ set of panels – see also Discussion).

Next, for the heterogeneity observed in red cones to usefully shape signal encoding in postsynaptic circuits, we reasoned that it must apply locally, including for immediately neighbouring cones. This would allow postsynaptic bipolar cell to integrate over a range of cones with differing sensitivities to ultimately encode a larger portion of the stimulus range. The dendrites of larval zebrafish bipolar cells a typically span some ∼20 µm in diameter^38^, which corresponds to ∼3-4 red cones in a row^39^. Accordingly, we next directly compared the dark-light index values (DLi) of neighbouring or small groups of red cones, which confirmed that heterogeneity in red cone output does occur locally. In an example line scan (Supplemental Figure S5A-E), neighbouring red cones within a 5 µm distance exhibit DLI values from -0.34 to -0.87. After plotting the difference in DLI as a function of distance between pairs of cones within the example line scan (Supplemental Figure S5E) and at the population level (Supplemental Figure S5F), we find the largest differences in DLI fall within 6 µm, with an average difference in DLI between neighbouring cones of 0.18. Similar local heterogeneity in DLI was consistently observed across most recordings (Supplemental Figure S5G).

### Variations in frequency-dependent output of individual cones

The decomposition of visual signals through retinal channels implementing different temporal filters is well-known at the level of retinal ganglion cells and has generally been thought to depend on inhibitory interactions between bipolar cells and amacrine cells in the inner retina^40–42^ but the results in Fig. 4 demonstrate that the synapses of individual cones can already be tuned differentially to emphasize either transient or sustained visual signals. The striking emphasis on the transient signalling of negative contrasts rather than positive at the output of red cones (Fig. 4E) may contribute to the observation that downstream retinal Off-circuits tend to transmit higher frequencies than On^43–45^. To investigate the temporal filters at red cone synapses we measured responses to a “chirp” stimulus in which the frequency of a full-field sinusoid (100% contrast) was swept from 20 Hz to 1 Hz over a period of 20 s. Fig. 5A shows two example cones with different responses to this chirp. Cone 1 tracked the stimulus throughout the frequency range, while cone 2 responded preferentially at lower frequencies. As before, different frequency-responses were routinely observed amongst neighbouring red cones recorded simultaneously. The frequency responses were characterized as the spectral centroid, the frequency where the centre of mass of the spectrum is located (Methods). Cones 1 and 2 had spectral centroids of 11.2 Hz and 7.5 Hz, respectively, and across 49 cones this central frequency ranged from ∼6 to 11 Hz (mean 9.2 ± 1.1) (Fig. 5B).

To assess the variability in the timing of responses, we next presented a 20 Hz sinusoidal stimulus at 100% contrast for 30 s (Fig. 5C). Red cones again responded heterogeneously: while some were nearly perfectly phase-locked to the stimulus (e.g. cone 3 in Fig. 5D) others appeared to respond more stochastically (cone 4). As a measure of the temporal precision, we calculated the vector strength^35^, a metric which varies from a value 1 for perfect synchronization to zero for random response timing. Cones 3 and 4 had a vector strength of 0.98 and 0.41, respectively. Across a population of 49 cones from 6 fish, the vector strength at 20 Hz ranged from 0.17 to 0.98 and averaged 0.61 ± 0.25 (± sd) (Fig. 5E). Together, the experiments summarised in Figs. 4,5 illustrate how ancestral red cones encode visual stimuli in a highly heterogeneous manner. The simple picture of a given cone type as representing a single temporal filter driving downstream circuits does not hold.

#### Heterogenous output of red cones depends on horizontal cell feedback

What is the source of this heterogeneity in the output of cone synapses? Do these variations reflect functional differences intrinsic to each cone or differences in the feedback signals from the network in which they are embedded? We reasoned that the relative contributions from these two possible sources might be disambiguated by pharmacologically isolating cones from the rest of the retinal circuit. This was achieved by blocking the cone drive to horizontal cells using the AMPA receptor antagonist CNQX injected into the eye (Methods, estimated final concentration: 50 µM)^6,46^, a manipulation expected to hyperpolarise horizontal cells and reduce their negative feedback to cones^15^.

A direct comparison of the output from one cone before and after inhibiting negative feedback is shown in Figs. 6A, where the stimulus is the same series of positive and negative contrast steps used in Fig. 4. With feedback intact, the sustained component of the response was approximately linear, but after blocking feedback the output was strongly rectified to negative contrasts. A similar pattern was observed in the contrast-response functions of 26 cones in 17 fish (Fig. 6B, where for simplicity the DLI is the sum of the transient and sustained release components). These results immediately suggest a possible mechanism for the varying degrees of rectification in the contrast-response functions of red cones: differences in the strength of negative feedback from horizontal cells adjusting the cone’s operational “set-point” (that is, baseline rates of vesicle release at this intermediate luminance).

**Figure 6.**
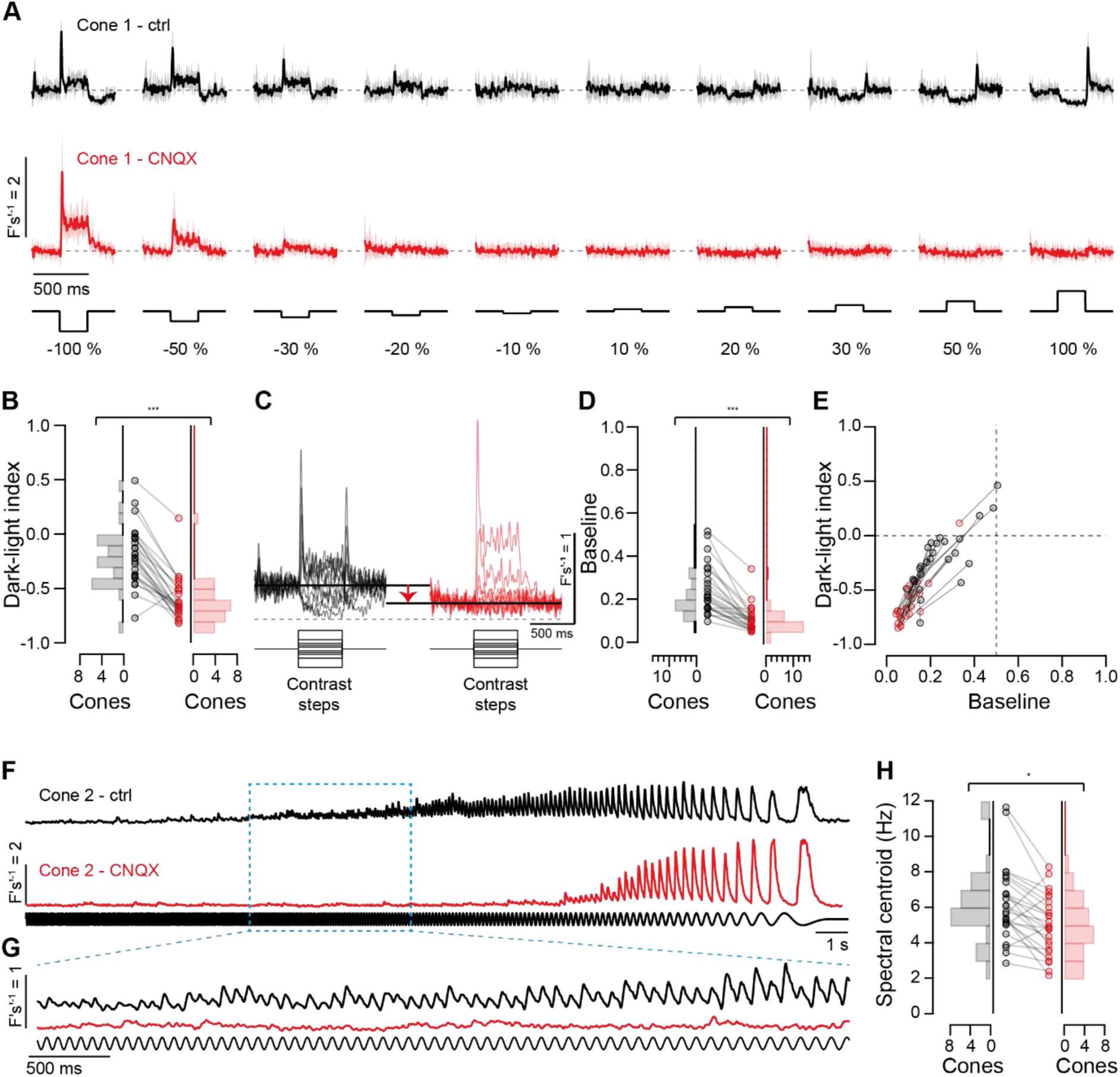
Cone heterogeneity requires intact outer retinal circuitry. **A**, Mean responses (± sd, shading) to contrast steps (cf. Fig. 4A) of one example cone before (black) and after (red) intraocular injection of CNQX to block horizontal cell feedback. **B**, This manipulation resulted in a systematic and significant drop in cone DLI values from -0.22 ± 0.27 to -0.63 ± 0.20 (p < 0.001, Wilcoxon rank sum, n = 26 cones in 17 fish) and value distribution (Chi-squared value = 16.1, critical value = 7.8, p < 0.001). **C**, Superposition of mean contrast responses from (A) to illustrate how dynamic range and baseline were extracted and **D**, comparison of baseline across experimental conditions; control: 0.24 ± 0.11, CNQX: 0.10 ± 0.06 (p < 0.001, Wilcoxon Rank Sum) and a significant decrease in heterogeneity (Chi-squared value = 8.8, critical value = 5.9, p < 0.05). **E**, Strong correlations between cone DLI and baseline value across both datasets. Control (black) Pearson correlation coefficient r = 0.77 (p < 0001); CNQX (red) r = 0.89 (p < 0.001), both datasets combined: r = 0.88 (p < 0.001). **F-H**, Glutamate release from an example red cone driven by chirp stimulus (cf. Fig. 5A) before (top, black) and after (red, bottom) CNQX injection (F) and expansion of the same data as indicated (G). Spectral centroids before (left, black) and after (right, red) CNQX injection, with a significant decrease (H, p < 0.05, n = 25 cones, 11 fish, Wilcoxon Rank Sum) but no significant change in heterogeneity (Chi-squared value = 7.3, critical value = 12.6, n.s.)

To explore this idea, we estimated each cones’ full coding range based on the minimum release rate at the highest light intensity and the maximum rate in darkness (Fig. 6C). We then measured the baseline as a fraction of this full range, which revealed a significant (*p* < 0.001, Wilcoxin Rank Sum) drop in set-point following block of feedback from horizontal cells (Fig. 6D). Crucially, the degree of rectification in cone output was strongly correlated with the baseline measured with or without block of negative feedback (Fig. 6E; Pearson’s correlation coefficient, control r = 0.77 (*p* < 0.001), CNQX r = 0.89 (*p* < 0.001). Along this tight relationship, pharmacological removal of horizontal cell feedback always shifted a cone’s behaviour towards a lower baseline and a correspondingly lower DLI. Blocking feedback caused both DLI and baseline values to become less heterogeneous across cones (Fig. 6C and E; variance of DLI values for control = 0.07, CNQX = 0.04, variance of baseline values for control = 0.01, CNQX = 0.003), with further significant decreases in the spread of DLI values.

The above observations were also largely mirrored in differences in cones’ temporal encoding in the presence and absence of feedback. When probed with the same chirp stimulus previously used to assess cones’ temporal encoding (Fig. 5), disconnecting horizontal cells shifted cone preferences to lower temporal frequencies (Fig. 6F-H; control = 6.2 ± 2.1 Hz, CNQX = 5.0 ± 1.7 Hz, Wilcoxon Rank Sum *p* < 0.05, n = 25 cones, 11 fish).

Together, these observations demonstrate that the set-point of a cone defines how it encodes positive versus negative contrasts and this set-point depends on the feedback received from horizontal cells. We conclude that a major source of heterogeneity in cone outputs are differences in their interactions with horizonal cells.

### Possible benefits of cone-heterogeneity for encoding natural contrasts

Variations in the input-output relation of the first synapse in vision is expected to impact all downstream processing. What might those impacts be? To explore this question, we set up a data-driven model of bipolar cells^47^ that sum cone inputs. We drove the model with naturalistic contrast series and compared responses to homogeneous and heterogeneous populations of cone synapses (Fig. 7). The results from this model indicate that heterogeneous sampling can improve representation of naturalistic contrast series (Fig. 7).

**Figure 7.**
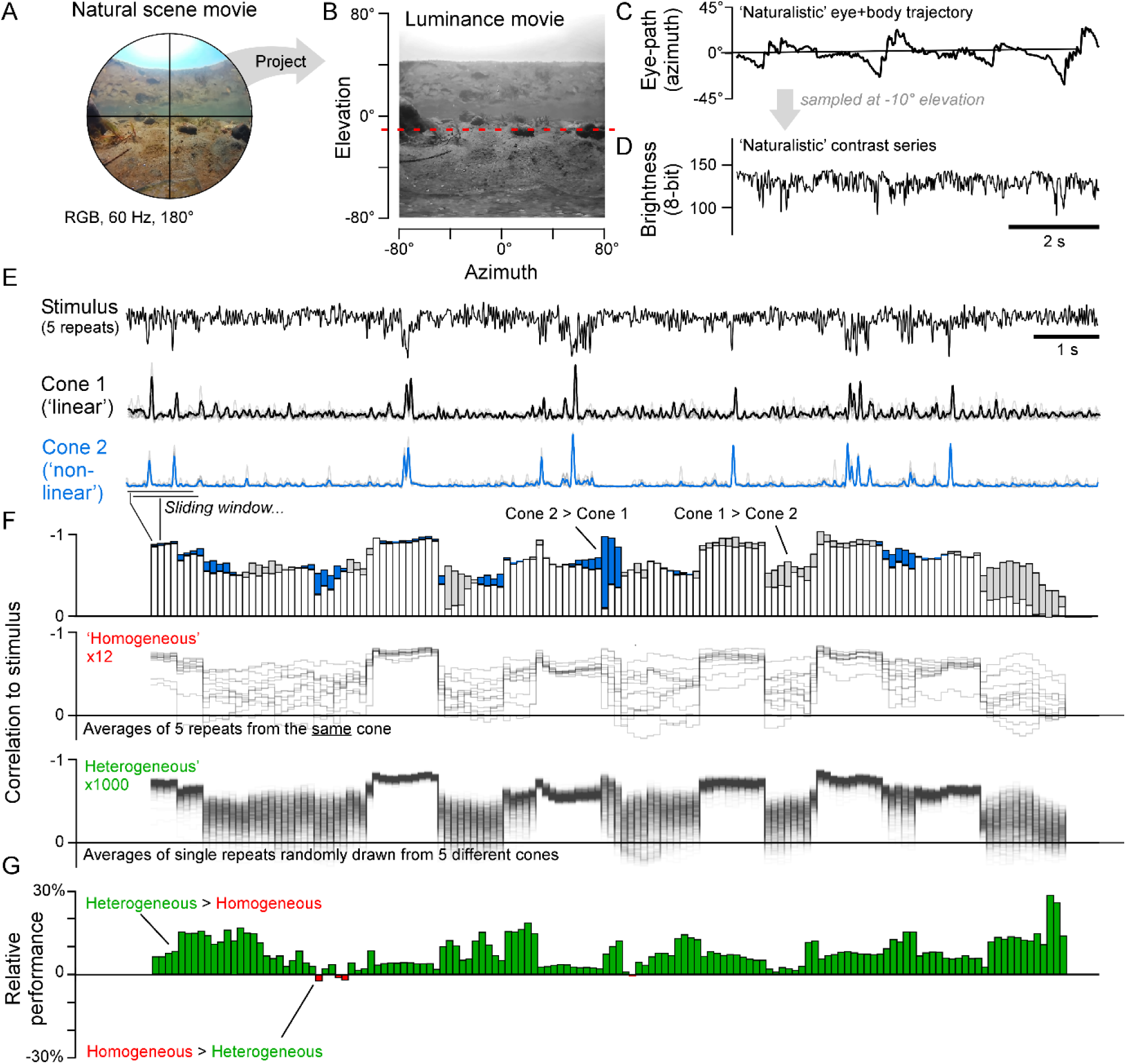
Data-driven model of heterogeneous versus homogeneous contrast encoding in natural scenes. **A-D**, A 180° fisheye natural scene movie recorded in India (A, from Ref^39^) was converted into a luminance signal over time projected onto a 2D plane (B, Methods). Next, a representative azimuth -only combined eye and body trajectory was extracted from a freely swimming zebrafish larvae (C, from Ref^48^) and mapped onto the natural scene move at a fixed elevation of -10° (dashed line in B) to extract a naturalistic contrast series (D). **E**, Responses of two example cones to the contrast sequence from (D) (mean superimposed on 5 repeats). **F**, top, Correlations between the example cone mean response and the stimulus, computed over a 1 second duration sliding window as indicated. Grey and blue filled areas highlight instances where the responses of cone 1 or cone 2, respectively, correlated more strongly with the stimulus. (F, middle), as top, each of the n = 12 cones recorded (“homogeneous encoding”), and (F, bottom), for 1,000 averages of 5 randomly sampled repeats across different cones (“heterogeneous encoding”). **G**, Comparison of the average performance of the “homogeneous” and “heterogeneous” populations of correlations from (F).

To arrive at this conclusion, we repeatedly stimulated individual red cones with a naturalistic contrast sequence extracted from the zebrafish natural environment (Fig. 7A-C). In the live eye, cones are challenged with ongoing patterns of light that vary in space, time, and intensity. However, from a cone’s perspective, and under the explicit caveat of leaving aside any added effects of cones’ centre-surround interactions (see discussion), changes in space as the eye sweeps across a visual scene can be mapped onto corresponding intensity changes over time. Accordingly, we combined available video data from the zebrafish natural environment^39^ (Fig. 7A,B) with data on eye- and body-movements from free-swimming zebrafish^48^ (Fig. 7C) to extract a 15 s “representative” naturalistic contrast series (Fig. 7D, Methods). We then presented five repeats of this contrast series to a total of n = 20 red cones from 8 fish.

As before, different red cones encoded this contrast sequence in a highly heterogeneous manner, here illustrated by the two example cones shown in Fig. 7E. Importantly, despite this heterogeneity *across* different cones, the responses of both cones *individually* were highly reproducible, reiterating cones’ exceptional response reliability and time precision (cf. Figs. 2,3). To explore how these two example cones differed in their encoding of this common contrast sequence, we next calculated each cone’s mean response across the five repeats and compared each to the original stimulus. We reasoned that the mean of five responses from a single cone is equivalent to the response of a hypothetical downstream neuron that averages individual responses from five functionally identical cones. In other words, the two mean responses mimic two different variants of a scenario in which the cone array is functionally homogeneous. To compare how accurately these two ‘homogenous’ cone-scenarios represented the contrast sequence, we calculated the correlation coefficient between each mean and the stimulus over a 1 s sliding window (Fig. 7F, top traces, Methods). Neither cone’s output was systematically superior to the other; during some phases of the sequence cone 1 correlated more strongly with the stimulus than cone 2, but in other phases this was reversed. It appears difficult to define an optimal set point for an individual cone.

Next, we tested bipolar cell responses with heterogenous cone inputs. Using the same five stimulus repeats from twelve recorded cones, we compared performance with i) a population of 12 bipolar cells, each summing across the five trials recorded from the *same* cone (as above; “homogeneous”), and ii) a population of 1,000 bipolar cells, each randomly summing any five trials from the full set of 60 trials available (“heterogeneous”). The individual performances of all modelled bipolar cells are shown in Fig. 7F middle/bottom. We then took the mean performance of either population as an indication of their central tendency and computed their relative differences (Fig. 7G, Methods) such that a difference of 0% indicates that both strategies, on average, perform equally well, while positive or negative percentages indicate correspondingly superior performance of the heterogeneous and homogeneous strategy, respectively. According to this simple comparison metric, the heterogeneous model systematically outperformed the homogeneous model by ∼8% on average, and by up to ∼30% depending on the detail in the stimulus. Moreover, the estimated performance boost was positive in >97% of time epochs lasting 1 s. This model provides a simple example of the way in which a functionally heterogeneous array of cone synapses may improve the coding of a naturalistic contrast series in neurons downstream.

## DISCUSSION

This study demonstrates that the synapses of red cones encode visual stimuli with exceptional reliability and time precision individually but are heterogeneous as a population in terms of sensitivity to luminance, contrast, and frequency (Figs. 1-5). These functional variations are tightly linked to differences in cones’ individual baseline “set-points”, which is in turn determined by interactions with horizontal cells in the outer retina (Figs. 6). A model of cone convergence on to bipolar cells indicates that this heterogeneity has the potential to improve the coding of naturalistic stimuli (Fig. 7). We also find that cones differentially encode different aspects of natural scenes via their transient versus sustained release components (Fig. 4). Together, these findings reveal a surprising degree of functional heterogeneity in the input to the retinal circuit and support the idea that photoreceptors interacting with their surrounding circuits can serve as different feature channels^17,18^ .

### Reliability of the cone synapse

Synapses inject noise into neural circuits because of the stochasticity of the processes that control vesicle fusion^49,50^ and in the retina this noise reduces the reliability of visual computations^30,51^. Once information is lost it can never be regained, so it is important that the first synapse in vision minimizes the noise introduced into the visual signal. Here we provide the first estimate of the reliability of the first synapse in vision *in vivo* and find that the variability is 10-fold less than expected for the Poisson process thought to operate at most synapses in the brain. The SNR of ∼90 per synapse for a 40 ms stimulus can be put into context by comparing it with the SNR of the *total* excitatory synaptic current that a mammalian alpha retinal ganglion cell receives from ∼500 bipolar cell inputs, which is ∼50-100^52,53^.

It has long been recognized that envisioning the ribbon synapses of rod photoreceptors as Poisson machines cannot easily account for the reliability with which single photons are detected and it has been suggested that a “clocking” mechanism might regularize the intervals between release of individual vesicles^54^. An alternative mechanism for making the synaptic output less variable at a given average release rate is the process of multivesicular release, where multiple vesicles are released as one synaptic event^55,56^. Electrophysiology in retinal wholemounts provides evidence for a combination of these mechanisms operating in rods under voltage-clamp stimulation^33^, although it is still not clear how they determine responses to light. The technique we have described here to isolate the output from individual cones has the advantage of providing *in vivo* measurements of glutamate release with a temporal resolution close to 1 kHz, which will allow the synaptic coding of visual information to be examined using, for instance, natural scenes. Crucially, it is now clear that ribbon synapses in rods, cones and bipolar cells do not encode visual information by modulating the rate of a Poisson process^26,33,57^. It remains to be seen how this improved reliability might depend on the specialized structure of ribbon synapses^36^ .

### Stimulus representation across the cone array

Sensory systems typically deal with two related types of mismatches between the statistical structure of the physical world and the encoding capability of sensory neurons: Incoming information tends to be linear and high bit-depth, but sensory neurons are usually nonlinear and have low bit-depth^58,59^. Consequently, populations of sensory neurons often use compromise solutions that disproportionately encode stimulus aspects that are most likely to be useful for the animal^60,61^. Here, our direct measurements of the light-driven cone drive to the retinal network *in vivo* offer new insights into the strategies used to meet those demands. We find that both the issue of limited dynamic range, and the issue of non-linear encoding, appear to be part-addressed already at this first synapse of vision.

#### Expanding coding range of natural contrasts

Strong spatial correlations in natural scenes^62^ mean that most of the time, neighbouring photoreceptors are driven by essentially the same stimulus over time. Moreover, downstream circuits tend to be driven by the simultaneous activity of more than one cone^63,64^. Accordingly, both from a sensory and a circuit perspective, cones do not operate in isolation. Instead, cones encode visual stimuli as a population, and we have shown that this population is locally heterogeneous (Fig. 1-4), potentially to expand the effective coding range of the visual system (Fig. 7). The presence of mechanistically distinct but conceptually related strategies in the early visual systems of mice^10^ and flies^12,65^ hints that locally heterogeneous sampling of the outside world may present a fundamental principle across convergently evolved visual systems. However, how possible improvements in overall signal representation scale with spatial scale, and in turn how this scale maps onto the operational spatial bandwidth of the system, remains important to explore in the future. Similarly, whether and how functional cone impacts more complex spatiotemporal receptive field properties of downstream retinal neurons, such as directionally selective circuits^66^, remains unclear.

Additionally, our observation that HC feedback facilitates red cone response speeds and the encoding of higher frequencies contrasts with our previous observation that UV cones instead become more sustained when HCs are blocked^6^. Similarly, in mice, blocking HCs increased low frequency signals ^67^. These findings suggest that kinetic interactions between HCs and photoreceptors are species and cone-type specific.

#### Light versus dark encoding

Natural visual contrasts tend to vary in two directions around an intermediate mean, but photoreceptors are fundamentally built to preferably represent one of these directions more readily than the other. This is because neurons in general, and synapses in particular, are subject to numerous nonlinearities^10,68–70^. In the context of cones, one particularly relevant type of nonlinearity occurs when their ribbon synapses are well-stocked^36^: In this case, a sudden rise in presynaptic calcium causes near instantaneous release of all vesicles from the readily releasable pool, and this leads to a sharp transient burst of release ^7,71,72^. However, the opposite is not the case: release does not cease in an equally transient manner if calcium suddenly drops, and such a hypothetical signal would also be more difficult to usefully read out by postsynaptic circuits. Consequently, a strong dark-bias in transient release (Fig. 4E,G) is probably inevitable for of most cones^7,10,73^. And yet, following this transient release, cones were notably more linear on average at the level of their sustained release (Fig. 4F,G), which is associated with the subsequent intermediate and reserve pools of vesicles^36^. In this way, a single cone can in effect “multiplex” two very different types of information, where large transient release events encode the presence of a high-contrast dark-transition, while slower-scale release modulations encode more nuanced light and dark transitions in a more balanced manner. Such a “compound code” could be readily read out by postsynaptic circuits, for example based on the kinetics of postsynaptic processes^5,47^.

The specific encoding of high-contrast dark-events by the transient release component is an example of highly pre-processed feature representation already at the first synapse of vision^17^. Rather than faithfully encoding detail in visual scenes – a representation that is achieved in parallel by the sustained component – the transient component will be disproportionately driven by the foreground alone. This is because underwater, contrast rapidly deteriorates with distance viewing distance^17,74^ . This underwater effect largely disappears in air^18^, but terrestrial species could use a similar strategy to disambiguate in-focus versus to out-of-focus visual structure^75^.

Notably, the disproportionate representation of large negative contrasts (as opposed to positive contrasts) may be an “accident of design” driven by the fact that vertebrate photoreceptors are Off cells (the same basic strategy would also work for On-photoreceptors disproportionately representing large positive contrasts), and may link with the observation that Off-circuits tend to disproportionately represent several elementary aspects of visual scene, including fast temporal contrasts and spectrally broad achromatic signals^44,45,76,77^. Accordingly, while various types of dark-biases in vertebrate retinal encoding have been linked to statistical dark -biases in some^37,78^ but not all^6,9,10,79^ natural scenes, it seems reasonable to include the intrinsic polarity bias of vertebrate photoreceptors as a key contributor.

#### Local and global heterogeneity

The local heterogeneity observed between the light responses of neighbouring red cones is largely driven by HC feedback. Red cones are contacted by H1-type HCs only. Two non-mutually exclusive mechanisms may underlie how H1 HCs differentially impact neighbouring cones to generate heterogenous responses to full-field light stimulation. First, possible functional cell-by-cell variability within the H1 population^14^ might locally propagate onto cones, if each is connected to a distinct subset of H1 neurons. Alternatively, or in addition, possible synapse-by-synapse variability at each HC-cone contact might impart functional heterogeneity onto the red-cone population. Beyond HCs, the persistence of a small degree of heterogeneity in the absence of HC inputs hints that additional factors may be important as well. Here, one possible source may lie in variations in the synaptic ultrastructure of cones themselves, for example with regards to the outer segment size and/or synaptic parameters (Supplemental Figure S1).

Beyond red cones, previous work showed that zebrafish UV cones are also functionally heterogeneous. However, unlike red, UV cones were “globally” heterogeneous but showed no obvious heterogeneity locally ^6,7^; These intriguing differences might relate to the very different roles served by the red and UV cone systems in zebrafish vision^17–19^. UV cones are approximately linear in the acute zone, but highly non-linear nasally.

### Spatial processing

In this work we used widefield stimuli modulated in time and intensity, but not space. Accordingly, in the future, it will be important to probe possible effects of cones’ known centre-surround structure^14,46,80^ on the encoding of spatially discontinuous pattens of light. Nevertheless, our widefield approach probes the most common naturalistic use case of a cone, where its centre and surround are driven concurrently. Here, our results from cones mirror those from previous work on bipolar cells, where like in cones, lateral interactions with inhibitory circuits serve to speed up, linearise and decorrelate visual feature representation at a population level^41^. Conversely, our recordings of cone activity in the pharmacological absence of horizontal cell feedback (Fig. 6) approximately mimic spatially restricted stimulation of the centre. In this case, our finding that cones become more non-linear and dark biased (Fig. 6B) implies that the same would happen to cones located near a spatial contrast edge. In the intact network, cones are therefore expected to encode the presence of spatial contrast in a highly dark biased manner – a putative effect that would then be exacerbated by the pre-existing dark-bias in cones’ transient release (Fig. 4G).

## METHODS

### Experimental Model

#### Animals

All procedures were carried out in accordance with the UK Animals (Scientific Procedures) act 1986, under the UK Home Office guidelines and approved by the animal welfare committee of the University of Sussex. Zebrafish larvae (*Danio rerio*) were housed in petri dishes (maximum 50 animals per dish) at 28°C under a standard 14:10 day/night light cycle (lights on at 8 am, lights off at 10 pm, with a 15-minute luminance ramp up or down accordingly). Animals were grown in E2 medium (Westerfield, 2000) with the addition of 200 μM 1-phenyl-2-thiourea (PTU, Sigma, P7629) from 1 day post fertilisation (dpf) to prevent further melanogenesis (Karlsson, Von Hofsten and Olsson, 2001).

The following previously published transgenic lines of zebrafish were used: *Tg(Cx55:5:nlsTrpR,-tUAS:SFiGluSnFR)* for expression of SFiGluSnFR in horizontal cells (Yoshimatsu *et al.*, 2021), *Tg(thrb:TdTomato*) for expression of TdTomato in red cones (Suzuki *et al.*, 2013), *casper* for minimised pigmentation (White et al., 2008) and *crystal* for minimised pigmentation and melanin (Antinucci and Hindges, 2016).

For all two-photon imaging experiments, zebrafish larvae were 6-7 dpf larvae. Imaging experiments were carried out in a temperature-controlled room (∼ 20°C) between 1 and 6 pm. Zebrafish larvae were immobilised in 2% low melting point agarose (Fisher Scientific, Cat: BP1360-100), positioned to lie on their side in an imaging chamber on top of a glass coverslip (thickness 0). The imaging chamber was filled with E2 fish medium, submerging the agarose-fixed larvae. Eye movements were further prevented by injection of α-bungarotoxin (1 nL of 2 mg/ml; Tocris, Cat: 2133) into the ocular muscles behind the eye.

#### Two-photon glutamate imaging

A Scientifica two-photon imaging system was used, equipped with a mode-locked Ti:Sapphire laser (Chameleon II, Coherent) tuned to 915 nm for excitation of the SFiGluSnFR reporter. Two-photon excitation was delivered through the objective (20X water-immersion, XLUMPlanFL, numerical aperture 0.95, Olympus). Emission of the SFiGluSnFR signal was captured above and below the sample, through the objective and condenser (Oil-immersion, numerical aperture 1.4, Olympus). The emitted signal was filtered through GFP filters (HQ 525/50, Chroma Technology) and detected by above- and sub-stage GaAsP photomultiplier tubes (PMTs, H7422P-40, Hamamatsu). The signal from the above- and sub-stage PMTs passed through a current-to-voltage converter before being summed with a custom-built summing amplifier and digitised. Image acquisition was controlled through ScanImage for Windows (3.8, MatLab). Functional recordings of the SFiGluSnFR signal were taken as 1 kHz line scans (128 X 1 pixels per frame, 1 ms per line). The dendritic tips of horizontal cells expressing SFiGluSnFR invaginate the photoreceptor pedicle (alongside bipolar cells (not labelled)) and act as an antenna for photoreceptor glutamate release. Line scans were positioned over the terminals of 3-5 cones per scan. Laser activation of photoreceptors was minimised by focusing the line scan across the pedicle, thus avoiding the photosensitive outer segment.

For consistency, all recordings were taken from the nasal retina, approximately aligned with the outward facing visual horizon (discussed in ^39^).

#### Light stimulation

Full-field visual stimulation was delivered from an amber light emitting diode (LED, 590nm) through a lightguide (both ThorLabs) positioned approximately 1 cm from the zebrafish retina. The light was filtered through a 590/10 nm bandpass filter (Thorlabs) to maximally activate red cone opsins, whilst minimally activating green cone opsins, and to fall outside of the activation range of blue and UV opsins. At full power, the output of the LED at the sample plane was ∼660 µW, sufficient to almost suppress glutamate release completely for tens of seconds. Stimulation protocols were driven through IGOR pro 6.3 for Mac (Wavemetrics), and the microscope was synchronised to visual stimulation.

#### Pre-processing

Movies were imported as tiff files in IGOR Pro 8.0 for Windows (Wavemetrics) and analysed using a suite of analysis routines; SARFIA^81^, GlueSniffer Analysis Package^26^ (James et al., 2019), and custom written procedures.

The GlueSniffer Analysis Package was used to analyse line scans and isolate SFiGluSnFR signals to individual cone terminals. Briefly, the spatial profile of a line scan was measured by making a temporal average of the fluorescence signal along the line scan. The average fluorescence profile was fit with a sum of Gaussians, where each Gaussian component corresponds to a ROI. For each ROI, the change in fluorescence over time was extracted, weighted towards the pixels in the centre of the spatial profile, facilitating significant denoising. The relative change in fluorescence, or DF/F, was calculated using the most frequent value (that is, the baseline of the recording). Any drift in the resulting DF/F trace was baseline corrected where required, using a linear correction. Recordings with strong photobleaching or low signal:noise were discarded. Additionally, recordings were discarded where an initial 100% contrast, 2 s off-step evoked a response with an amplitude of less than 3 standard deviations of the baseline.

The possibility of conflating SFiGluSnFR signals from multiple cone terminals was limited experimentally by taking line scans at spatially separated ROIs, avoiding planes of focus where multiple ROIs were crowded together. Second, the small point-spread function (PSF) of the microscope reduced conflation of signals. The microscope used for these experiments has a PSF of 0.7 μm in x and y, meaning the synapses of cones >1 μm apart could be easily distinguished. The PSF in z was 2.2. μm. The average cone terminal width was approximately 2-3 μm (Supplemental Figure S1). Moreover, during imaging the centre of the cone terminals of interest were approximately identified by the strongest SFiGluSnFR signal, and line scans were taken at this imaging plane to minimise the conflation of signal from terminals above and below the plane of focus was minimised.

Next, DF/F traces of the SFiGluSnFR signal were deconvolved to limit the decay element of the SFiGluSnFR fluorescent reporter from the recordings and produce traces that are proportional to the rate of glutamate release, or DF/F per second (F’s^−1^). This was achieved with a Wiener filter, as described previously^26^. The decay of the SFiGluSnFR reporter was estimated by fitting transient responses with a kernel. Transients at most cone terminals could be described with a kernel with a decay of 0.06 s. By filtering the decay of the fluorescent reporter from the signal, it was possible to recover an estimate of the glutamate signal.

#### Dark-light index

The dark-light index (DLI) was calculated from the cone’s contrast-response function; the area under the curve (AUC) for the negative contrast steps was measured, giving value “b”, and the AUC for the positive contrast steps was measured, giving value “a”. To calculate an index value, both numbers must be positive, therefore the “a” value was multiplied by -1. Following this, if the “a” value was negative (i.e., because the contrast response curve sat above zero for the positive contrast steps due to a noisy signal), the “a” value was assumed to be zero. The dark-light index was calculated as (a-b)/(a+b), producing a DLI value between -1 and 1. Where the DLI = -1, the cone exclusively responded to negative contrast steps, where the DLI = 0 the cone responded to both positive and negative contrast steps equally, and where DLI = 1 the cone responded exclusively to positive contrast steps.

#### Spectral centroid

Glutamate responses to the 20 Hz chirp stimulus were analysed to identify the spectral centroid, that is, the “centre of mass” of the frequencies in the glutamate responses, following:

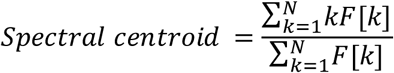

Briefly, the Fourier transform (FT) of the response trace was weighted by the power of the frequencies. The sum of this weighted FT was divided by the sum of the FT to produce the spectral centroid.

#### Vector strength

Glutamate responses to the 20 Hz sinusoid stimulus were analysed to measure the phase locking of responses over repeats of the stimulus phase. Responses were detected by differentiating the glutamate records and using the in-built peak finding function in Igor Pro 8.0 for Windows, “PeakFinder”, to detect the rise of the response transient. The mean + 1 sd of the glutamate signal during exposure to mean-light levels (between 6 and 8 s of stimulus) was applied as the threshold for detecting a response. Response times were converted to a phase angle, θ_*i*_, i.e. where they occur within a stimulus period. Briefly, where one stimulus phase is equal to 360 degrees, the time of the rise of each response was converted to the degree at which it occurred in the stimulus phase, and then to radians. The vector strength was calculated, as described elsewhere^82^, as follows:

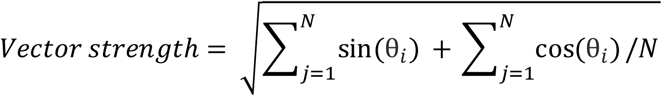

The vector strength for each cone recording is the square root of the sum of the sine and cosine of the vector values for each response. The vector strength may vary from 0 to 1, with 1 implying perfect synchronisation of responses with the stimulus.

#### Statistics

Statistical analysis was carried out using in-built functions of IGOR Pro 8.0 for Windows (Wavemetrics). When data were not normally distributed, nonparametric tests were applied. Tests were two-sided and significance was defined as *p* < 0.05. Where required, post-hoc tests were used to correct for multiple groups. If an experiment required the delivery of more than one visual stimulus protocol, the order of the stimuli was randomized. Data collection and analyses were not carried out blind as red cone responses were specifically selected for during experiments and analysis. Recordings were excluded from analysis if a low signal to noise ratio of the deconvolved SFiGluSnFR signal impeded accurate detection of light responses to off-steps of light.

#### Pharmacology

For some experiments, HCs light responses were blocked with an injection of cyanquixaline (CNQX, Tocris, Cat: 1045, final concentration of approximately 50 µM) in artificial cerebro-spinal fluid (aCSF). CNQX is a synthesized non-NMDA (N-Methyl-D-Aspartate) receptor antagonist that blocks AMPA- and kainate-receptors, effectively blocking HC light responses. CNQX was injected intravitreally into the eye, through the cornea adjacent to the lens (usually temporal to the lens).

Beyond blocking cone input to HCs, CNQX additionally affects downstream retinal circuitry, which in turn could modulate the neuromodulator milieu of cones, for example via dopaminergic interplexiform cells (DA-IPCs)^83,84^. However, this is unlikely to cause major to off-target effects on red cones and associated synaptic pathways, becasue (i) DA-IPCs are thought to be primarily driven by the brain and (ii)previous work points to the rod system, rather than cones, as the major target for dopaminergic modulation in the zebrafish retina^85^.

#### Immunohistochemistry and confocal imaging

The transgenic line expressing TdTomato in red cones (*Tg(thrb:TdTomato)*) was outcrossed with the *crystal* line, and the resulting larvae were screened for TdTomato expression and a *crystal* phenotype (clear, non-pigmented eyes). The positive larvae were euthanised by tricaine methanesulfonate (MS222, Sigma Aldrich) overdose and fixed in 4% paraformaldehyde (PFA, Agar Scientific, AGR1026) in 1 X PBS on a rocker for 30 minutes exactly at room temperature. After three washes in 1 X PBS, whole eyes were enucleated, and the cornea was removed by using the tip of a 30 G needle. The dissected eyes were incubated in 0.1% Triton X-100 (Sigma, X100) made up in 1 x PBS for 15 minutes at room temperature. In fresh 0.1% Triton X-100, primary antibody was added; anti-1D4 (Santa Cruz, mouse, sc-57432) at 1:20. Samples were incubated at 4°C for 4 days. Samples were washed five times in 0.1% TritonX-100 in 1X PBS and treated with DAPI nuclear dye (Invitrogen, 33342) at 1:2000, and secondary antibody; Donkey-anti-mouse Dylight 647 (ThermoFisher, A32787) at 1:200. After one day incubation at 4°C, samples were washed in 0.1% Triton X-100 in 1X PBS twice. Samples were mounted in 1.5% agarose in PBS on a coverslip. The eyes were manually oriented with the lens facing down, and the PBS was replaced with mounting media (Vectorlabs, Vectashield, H-1000) for imaging.

Confocal image stacks were taken on an LSM880 (Zeiss) using a 40X oil immersion objective (plan apochromat 40X/1.3 oil DIC UV-IR M27), or a 63x oil immersion objective (HC PL APO CS2, Leica). Contrast, brightness, and pseudo-colour were adjusted for display in Fiji (NIH). Quantification of soma length, outer segment length, and terminal width were performed using custom scripts in IGOR Pro 8.0 (Wavemetrics) after manually marking the inner and outer locations of each organelle of interest.

#### Electron microscopy and analysis

Serial blockface scanning electron microscopy (SEM) volumes were gratefully received from Prof Rachel Wong (University of Washington, USA), as previously part-analysed and published in Ref^14^. The data consist of 652 vertical slices through the OPL of the Acute Zone region of the retina of a 6 dpf wild-type larval zebrafish. There is 50 nm between each layer, and the xy pixel dimension is 5 nm.

Using TrakEM2 plugin got FIJI, the data set was manually aligned and structures of interest (e.g., photoreceptor outer segment, mitochondria, nucleus, terminal, and HC and BC nuclei and dendritic projections) were manually traced. From here, 3D reconstructions of the traced structures were generated, and basic measurements e.g. volumes and surface areas, were calculated using in-built functions of FIJI.

### Modelling bipolar cell responses

A “representative” stimulus segment was extracted by combining a previously published^39^ video-segment (60 Hz) showing an underwater scene of a typical zebrafish natural habitat in Northern India with a “typical” azimuth-only eye and body movement trajectory measured in a free swimming zebrafish^48^ as shown in Fig. 7A-D. To this end, the video was converted to 8-bit greyscale by averaging the RGB components, followed by reading out of the single pixel brightness values over time (1 pixel per frame) as dictated by the eye-body movement trajectory. For implicity we did not include eye movements in elevation and kept the entire sequence at -10 degrees relative to the visual horizon, approximately aligned with the position that would be sampled by the nasal red cones surveyed during physiology. We presented five repeats of this stimulus sequence as a 60 Hz widefield stimulus to a total of 20 cones and extracted their glutamate responses. Twelve out of these 20 cones passed a quality criterion of 0.4 (see definitions in Ref^86^) and were used for the presented analysis.

## Acknowledgements

We thank João Marques and Mike Orger for sharing pre-publication tracking data from free-swimming zebrafish

## Author contributions

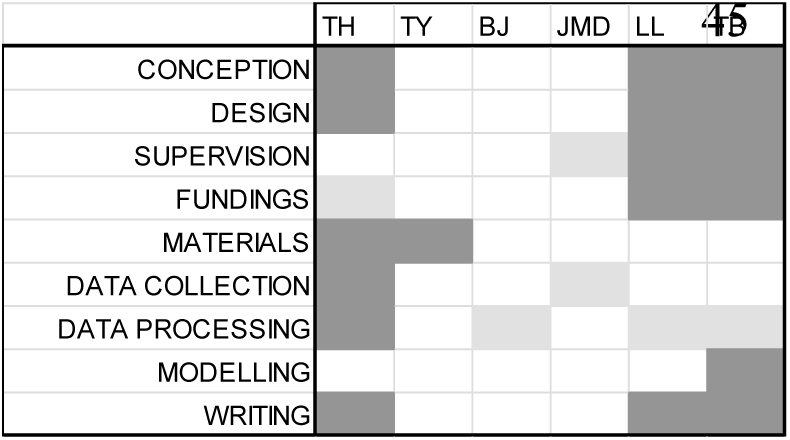

## Supplementary Figures

**Supplemental Figure S1.**
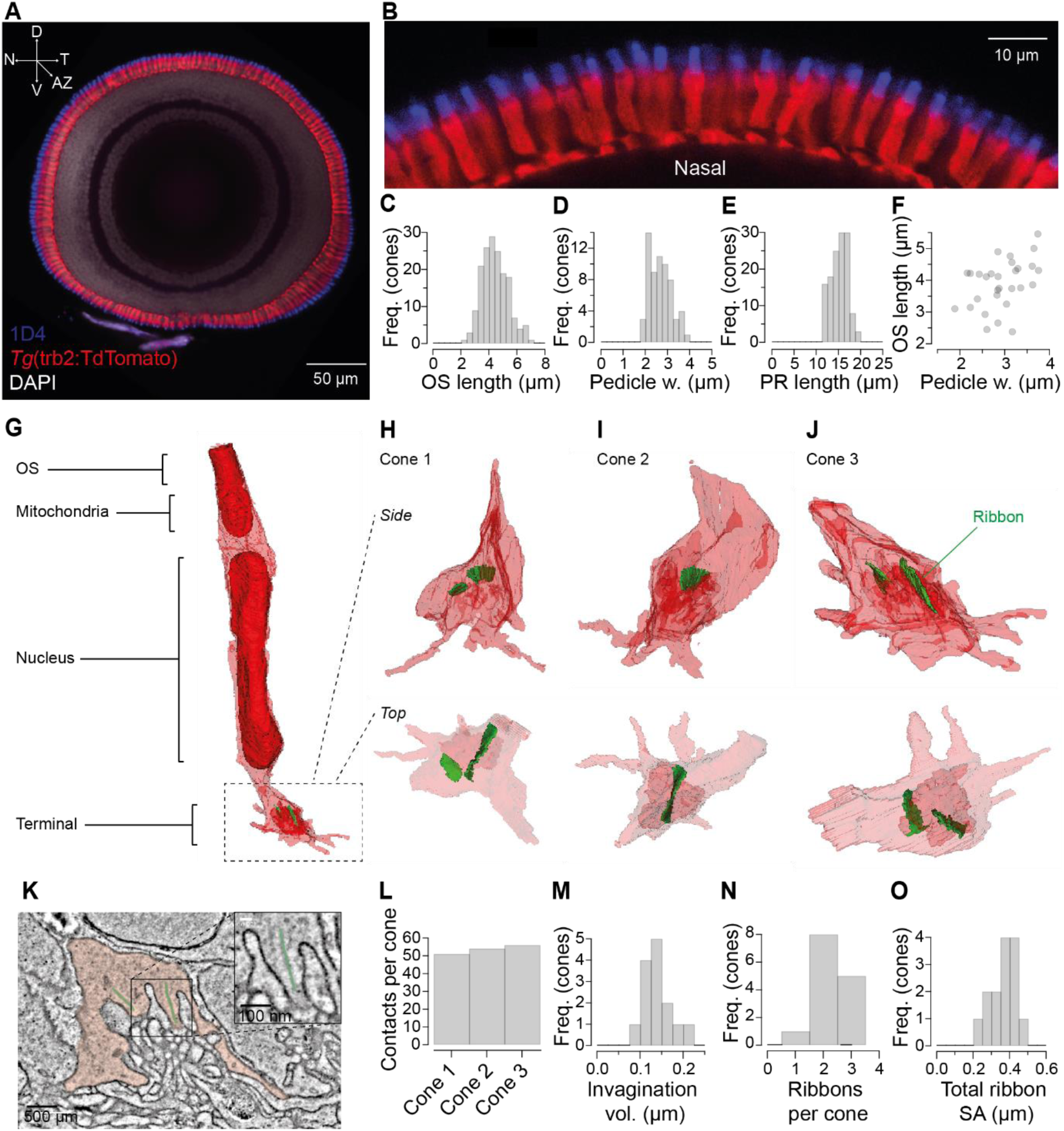
**A,** A whole-eye immunostaining of a 6dpf Tg(trb2:TdTomato) larvae (red cones, red), labelled against 1D4 (red cone outer segments, blue) and DAPI (nuclear stain, grey). D: dorsal; N: nasal; V: ventral; AZ; acute zone. **B,** An expansion of the nasal region from the same sagittal section. **C-E,** OS length, pedicle width and PR length plotted as histogram. **F,** OS length is plotted against pedicle width, demonstrating a positive correlation (Pearson correlation co-efficient r = 0.43, p = n.s.). **G,** A 3D-reconstruction of an example red cone (cone 1) from the acute zone region of a 6 dpf larval zebrafish. Dark red: outer segment (OS), mitochondria and nucleus, green: ribbons. **H-J,** Zoom-ins of 3 example red cone terminals with ribbons in green, shown from the side (top row) and from above (bottom row). **K,** An electron micrograph of a vertical section through the OPL, showing the terminal of cone 1 (transparent red) and its ribbons (transparent green). Insert shows a zoom in of two HC dendrites flanking a ribbon (transparent green). **L** The number of invaginating contacts per cone terminal for cones 1 – 3 (from H-J), including HC and BPC dendrites and cone telodendrites. Histograms showing **M,** the volume of the red cone terminal invagination, **N,** the number of ribbons per cone terminal and **O,** total ribbon surface area per cone terminal. For C n =168 red cones, 6 retinas, D n = 63 red cones, 5 retinas, E 130 red cones, 6 retinas, F 29 red cones, 4 retinas, M-O 14 cones, 1 retina.

**Supplemental Figure S2.**
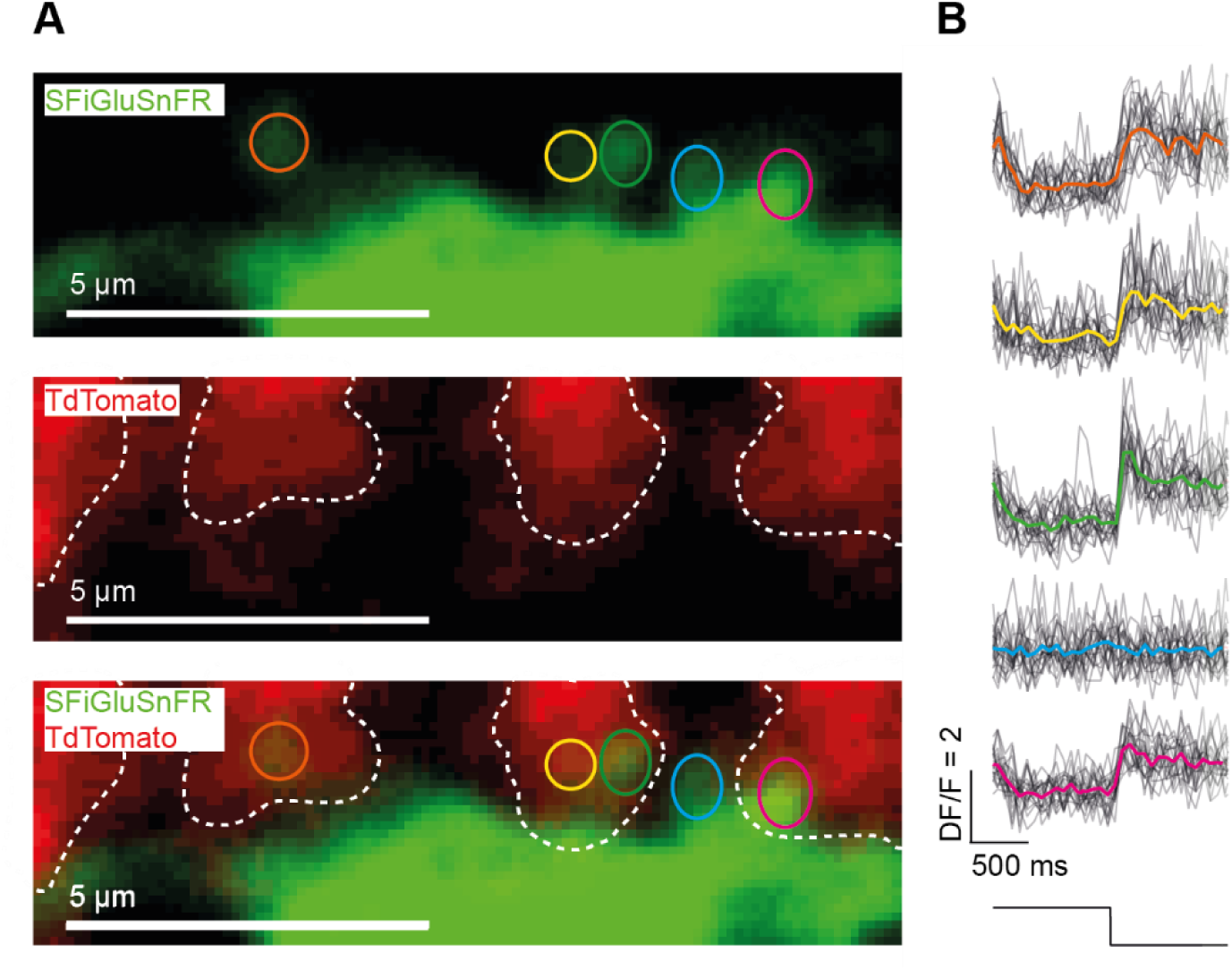
**A,** Maximum intensity projections of the field of view from a two-photon recording, focused on the OPL of a 6 dpf larval zebrafish, showing the expression of SFiGluSnFR in HCs (top panel), TdTomato in the terminals of red cones (middle panel), and an overlay of the two channels (bottom panel). The HC dendritic projections are circled as ROIs (top), and the red cone terminals are outlined with a white dashed line (middle). The overlay (bottom) demonstrates the blue ROI does not contact a red cone. **B**, SFiGluSnFR responses at each ROI in response to 2 s steps of light from full brightness to darkness. The on-off step stimulus was repeated 15 times. Responses to each trial are overlaid (black) and the mean is super-positioned (colour corresponding to ROI in (A)).

**Supplemental Figure S3.**
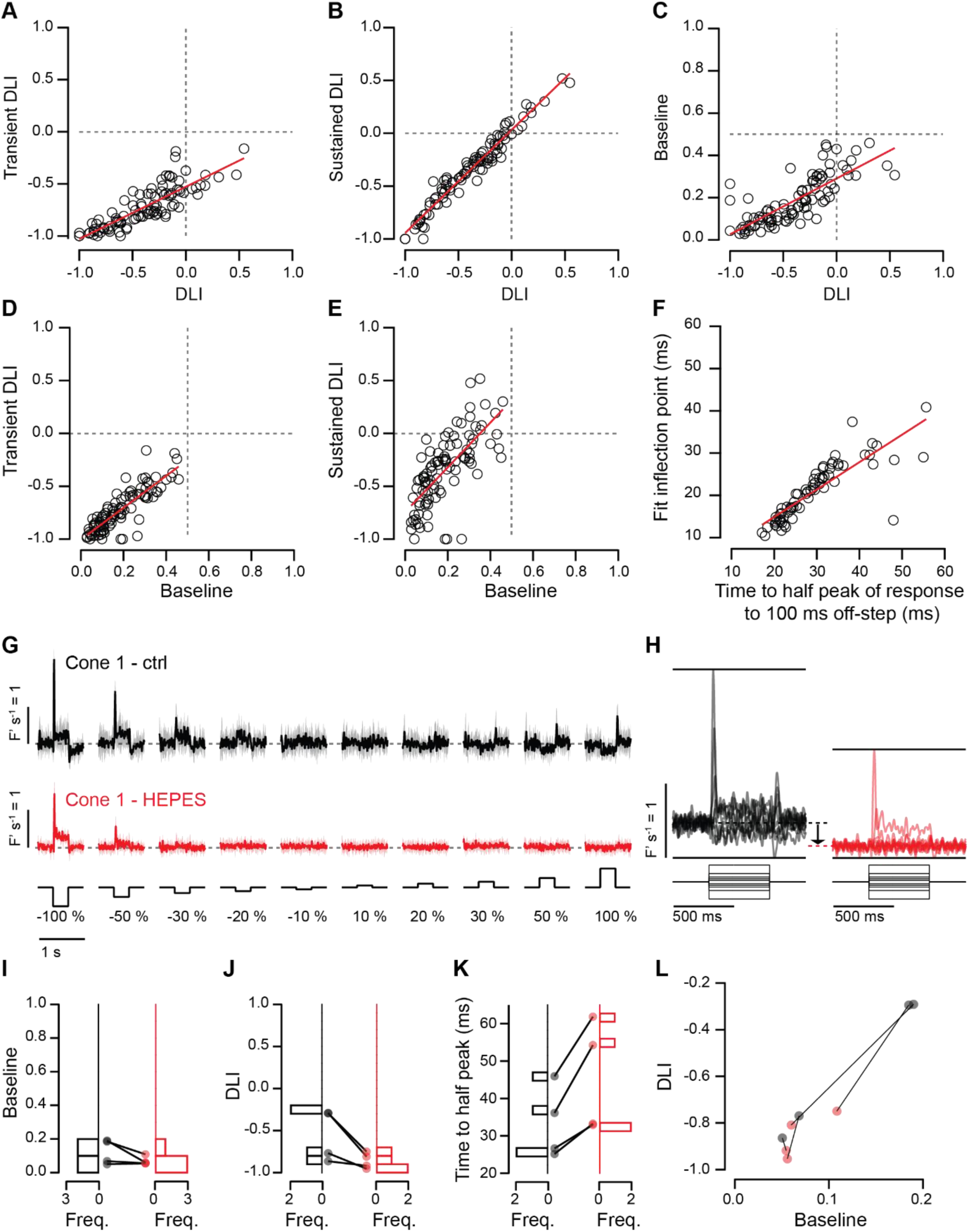
Correlation in response metrics. Strong, positive correlation was found in metrics measured in red cone glutamate responses to luminance contrast. **A**, Transient DLI and dark light-index (DLI) (r = 0.85, p < 0.0001), **B** sustained DLI and DLI (r = 0.98, p < 0.001), **C** baseline and DLI (r = 0.79, p < 0.0001), **D** transient DLI and baseline (r = 0.86, P < 0.0001), and **E** sustained DLI and baseline (r = 0.71, p < 0.0001). **F** Strong, positive correlation was found in red cone glutamate responses to pseudorandomised 5 – 100 ms off-steps, when considering the speed of responses to the 100 ms condition and the most sensitively encoded off-step durations (r = 0.84, p < 0.0001). **G** Contrast responses of an example cone before (top, black) and after intraocular HEPES injection (bottom, red). **H** For the same example cone, the mean responses to all stimulus conditions are superimposed, to demonstrate a drop in baseline after HEPES injection (dashed black and dashed red lines, arrow). **I – J** Baseline and DLI exhibit a reduction after HEPES injection, and **K** time to half peak of responses for the -100% contrast condition exhibit an increase after HEPES injection. **L** DLI is plotted as a function of baseline before (black) and after (red) HEPES injection, with grey lines linking the values for a single cone. (A-E) n = 97 red cones, 30 fish, (F) n = 70 red cones, 12 fish, and (I-L) n = 4 red cones, 2 fish.

**Supplemental Figure S4.**
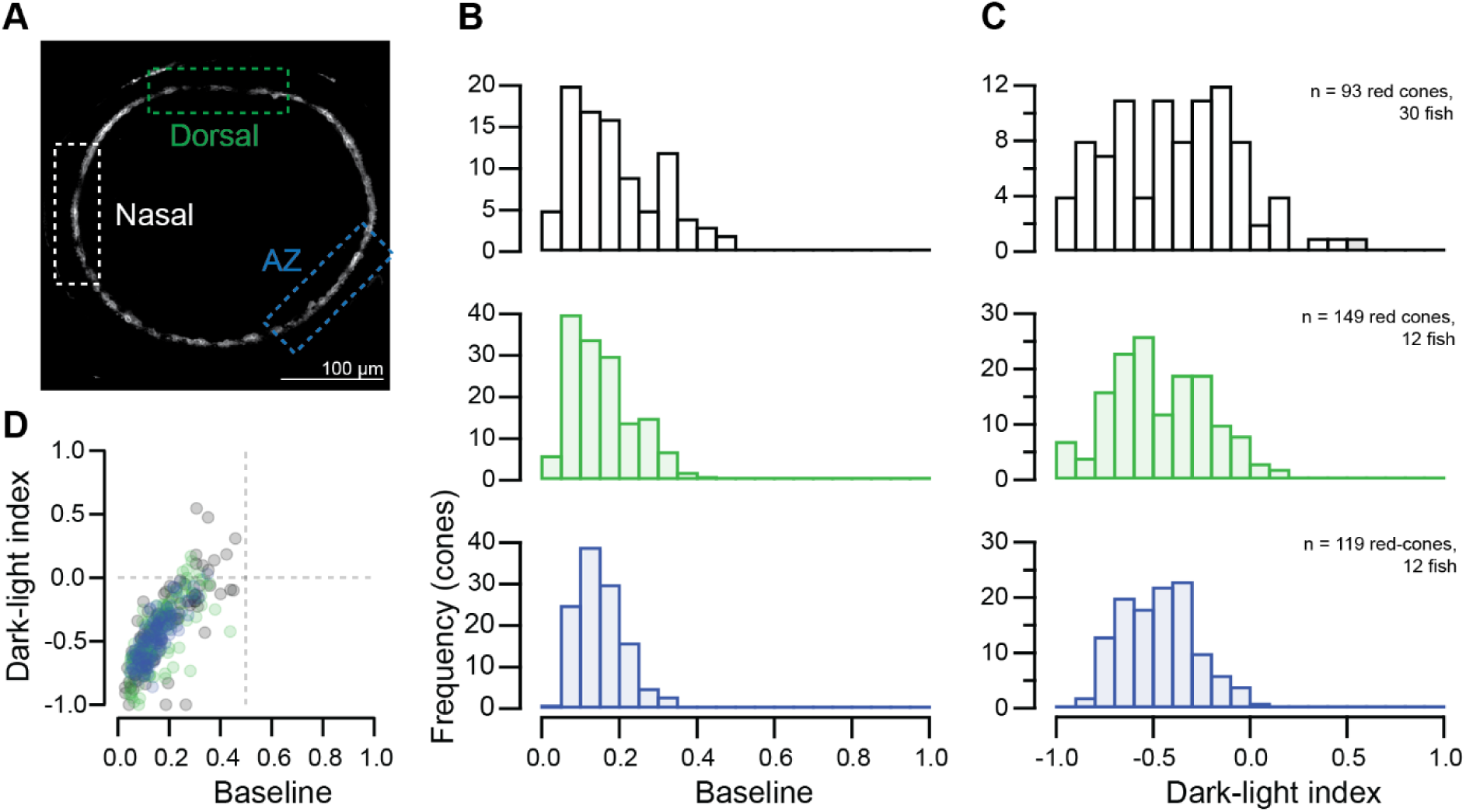
Heterogeneity in contrast responses present across retinal regions. **A**, Nasal (white), dorsal (green), and acute zone (AZ, blue) retinal regions are demarcated by dashed-line boxed superimposed on a two-photon image of the retina of a 6 dpf larval zebrafish expressing SF-iGluSnFR in horizontal cells (white). Glutamate responses to luminance contrast steps (see Fig. 4A) in nasal, dorsal and AZ retinal regions were recorded and the distributions of baseline, **B**, and dark-light index (DLI), **C**, are plotted as histograms for each retinal region (from top to bottom; nasal, dorsal, AZ). There is not statistically significant difference in the distribution of baseline nor DLI values between retinal regions (Kruskall-Wallace test for baseline H (2) = 4.34, p = 0.11, and for DI H(2) = 4.34, p = 0.11). **D**, The relationship between DLI and baseline for each cone is plotted, colour coded by retinal region. Nasal, n = 93 red cones, 30 fish, dorsal, n = 149 red cones, 12 fish, and AZ, n = 199 red cones, 12 fish.

**Supplemental Figure S5.**
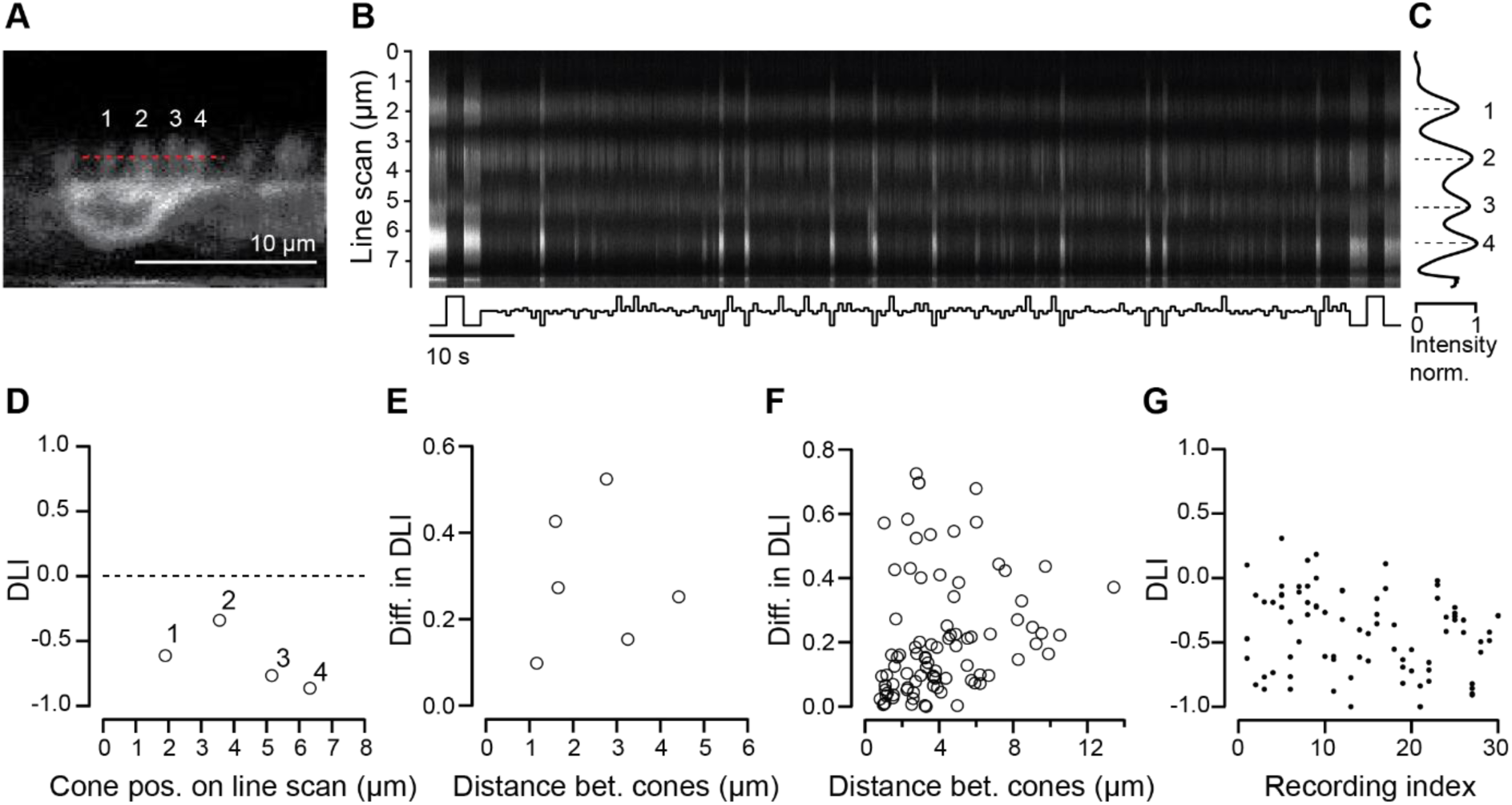
Local heterogeneity in cone contrast responses. **A**, Two-photon image of SFiGluSnFR expressed in HCs in the nasal retina of a 6 dpf zebrafish. The red dashed line marks the approximate position of the line scan across the dendritic processes of 4 cones. **B**, A kymograph of the line scan shows how fluorescence changes in response to full-field stimulation with contrast steps (bottom). **C**, The fluorescence profile of the line scan shows 4 spatially separated ROIs. The position of peak fluorescence (dashed lines) is taken as the position of each cone of the line scan. **D,** DLI is plotted as a function of cone position on the line scan. **E**, The difference in DLI values between each cone is plotted as a function of distance between cones for the above example, and for the population in **F**. **G**, DLI values per animal (recording index) are ordered from largest to smallest spread. (F – G) n = 84 red cones, 27 fish.

